# γδ T cells are effectors of immune checkpoint blockade in mismatch repair-deficient colon cancers with antigen presentation defects

**DOI:** 10.1101/2021.10.14.464229

**Authors:** Natasja L. de Vries, Joris van de Haar, Vivien Veninga, Myriam Chalabi, Marieke E. Ijsselsteijn, Manon van der Ploeg, Jitske van den Bulk, Dina Ruano, John B. Haanen, Ton N. Schumacher, Lodewyk F.A. Wessels, Frits Koning, Noel F.C.C. de Miranda, Emile E. Voest

**Affiliations:** Department of Pathology, Leiden University Medical Center, Leiden, the Netherlands; Department of Immunology, Leiden University Medical Center, Leiden, the Netherlands; Department of Molecular Oncology and Immunology, Netherlands Cancer Institute, Amsterdam, The Netherlands; Oncode Institute, Utrecht, the Netherlands; Division of Molecular Carcinogenesis, Netherlands Cancer Institute, Amsterdam, the Netherlands; Gastrointestinal Oncology, Netherlands Cancer Institute, Amsterdam, the Netherlands; Medical Oncology, Netherlands Cancer Institute, Amsterdam, the Netherlands; Faculty of EEMCS, Delft University of Technology, Delft, the Netherlands

**Keywords:** Mismatch-repair deficiency, Colon cancer, Immune checkpoint blockade, γδ T cells, β2-microglobulin (β2m), Human Leukocyte Antigen class I (HLA class I)

## Abstract

DNA mismatch repair deficient (MMR-d) cancers present an abundance of neoantigens that likely underlies their exceptional responsiveness to immune checkpoint blockade (ICB)^1,2^. However, MMR-d colon cancers that evade CD8^+^ T cells through loss of Human Leukocyte Antigen (HLA) class I-mediated antigen presentation^3–6^, frequently remain responsive to ICB^7^ suggesting the involvement of other immune effector cells. Here, we demonstrate that HLA class I-negative MMR-d cancers are highly infiltrated by γδ T cells. These γδ T cells are mainly composed of Vδ1 and Vδ3 subsets, and express high levels of PD-1, activation markers including cytotoxic molecules, and a broad repertoire of killer-cell immunoglobulin-like receptors (KIRs). *In vitro*, PD-1^+^ γδ T cells, isolated from MMR-d colon cancers, exhibited a cytolytic response towards HLA class I-negative MMR-d colon cancer cell lines and β*2*-*microglobulin* (*B2M*)-knockout patient-derived tumor organoids (PDTOs), which was enhanced as compared to antigen presentation-proficient cells. This response was diminished after blocking the interaction between NKG2D and its ligands. By comparing paired tumor samples of MMR-d colorectal cancer patients obtained before and after dual PD-1 and CTLA-4 blockade, we found that ICB profoundly increased the intratumoral frequency of γδ T cells in HLA class I-negative cancers. Taken together, these data indicate that γδ T cells contribute to the response to ICB therapy in patients with HLA class I-negative, MMR-d colon cancers, and illustrate the potential of γδ T cells in cancer immunotherapy.

## Introduction

Immune-checkpoint blockade (ICB) targeting the PD-1/PD-L1 and/or CTLA-4 axis provides durable clinical benefit to patients with DNA mismatch repair-deficient (MMR-d)/Microsatellite Instability-High (MSI-H) cancers^8–11^. The exceptional responses of MMR-d/MSI-H cancers to ICB are likely explained by their vast burden of putative neoantigens, which originate from the extensive accumulation of mutations in their genomes^1,2^. This is in line with the current view that PD-1 blockade mainly boosts endogenous antitumor immunity driven by CD8^+^ T cells, which recognize Human Leukocyte Antigen (HLA) class I-bound neoepitopes on cancer cells^12–14^. However, MMR-d colon cancers frequently lose HLA class I-mediated antigen presentation due to silencing of HLA class I genes, inactivating mutations in β*2-microglobulin* (B2M), or other defects in the antigen processing machinery^3–6^, which may render these tumors resistant to CD8^+^ T cell-mediated immunity. Interestingly, the majority of β2m-deficient MMR-d cancers have shown durable responses to PD-1 blockade^7^, suggesting that immune cell subsets other than CD8^+^ T cells contribute to these responses.

Immune cell subsets capable of HLA class I-independent tumor killing include natural killer (NK) cells and γδ T cells. γδ T cells share many characteristics with their αβ T cell counterpart, such as cytotoxic effector functions, but express a distinct TCR composed of a γ and δ chain. Different subsets of γδ T cells are defined by their TCR δ chain usage, of which those expressing Vδ1 and Vδ3 are primarily “tissue-resident” at mucosal sites, whereas those expressing Vδ2 are mainly found in blood^15^. Both adaptive and innate mechanisms of activation, e.g., through stimulation of their γδ TCR or innate receptors such as NKG2D, DNAM-1, NKp30 or NKp44, have been described for γδ T cells^16^. Killer-cell immunoglobulin-like receptors (KIRs) are expressed by γδ T cells and regulate their activity depending on HLA class I expression^17^. Furthermore, γδ T cells were found to express high levels of PD-1 in MMR-d colorectal cancers (CRCs), suggesting that these cells may be targeted by PD-1 blockade^18^.

Here, we applied a combination of transcriptomic and imaging approaches for an in-depth analysis of ICB-naïve and ICB-treated MMR-d colon cancers, as well as *in vitro* functional assays, and found evidence indicating that γδ T cells mediate responses to HLA class I-negative, MMR-d tumors during ICB therapy.

## Results

### γδ T cells are enriched in *B2M*-mutant MMR-d cancers

To gain insights into immune cell subsets involved in immune responses towards HLA class I-negative MMR-d cancers, we studied the transcriptomic changes associated with genomic loss of *B2M* in three MMR-d cancer cohorts in The Cancer Genome Atlas (TCGA): colon adenocarcinoma (COAD; n=44 *B2M^WT^*, n=6 *B2M^MUT^*), stomach adenocarcinoma (STAD; n=48 *B2M^WT^*, n=13 *B2M^MUT^*), and endometrium carcinoma (UCEC; n=99 *B2M^WT^*, n=3 *B2M^MUT^*). We found that *B2M* was among the most significantly downregulated genes in *B2M^MUT^* tumors (two-sided P=8.4×10^-5^, Benjamini-Hochberg corrected false discovery rate [FDR]=0.040; Fig. 1a). Genes encoding components of the HLA class I antigen presentation machinery other than β2m were highly upregulated in *B2M^MUT^* tumors (Fig. 1a). Interestingly, we found *TRDV1* and *TRDV3*, which encode the variable regions of the δ1 and δ3 chains of the γδ T cell receptor (TCR), among the most significantly upregulated loci in *B2M^MUT^* tumors (*TRDV1*: FDR=0.0056; *TRDV3*: FDR=0.0062; Fig. 1a). In line with this, the expression level of γδ TCRs was significantly higher in *B2M^MUT^* compared to *B2M^WT^* MMR-d cancers (Wilcoxon rank sum-based two-sided P=1.1×10^-5^ for all cohorts combined; Fig. 1b), while this was less pronounced for αβ TCR expression (Wilcoxon rank sum-based two-sided P=0.023 for all cohorts combined; Extended Data Fig. 1). In addition, multiple KIRs showed clear overexpression in *B2M^MUT^* tumors (Fig. 1a), where the expression level of all human KIRs combined was significantly higher in *B2M^MUT^* compared to *B2M^WT^* MMR-d tumors (Wilcoxon rank sum-based two-sided P=3.0×10^-7^ for all cohorts combined; Fig. 1c). Together, these results suggest that ICB-naïve, *B2M^MUT^* MMR-d cancers show increased levels of (1) Vδ1 and Vδ3 T cells and (2) immune cells expressing KIRs, receptors implied in the recognition and killing of HLA class I-negative cells.

**Figure 1.**
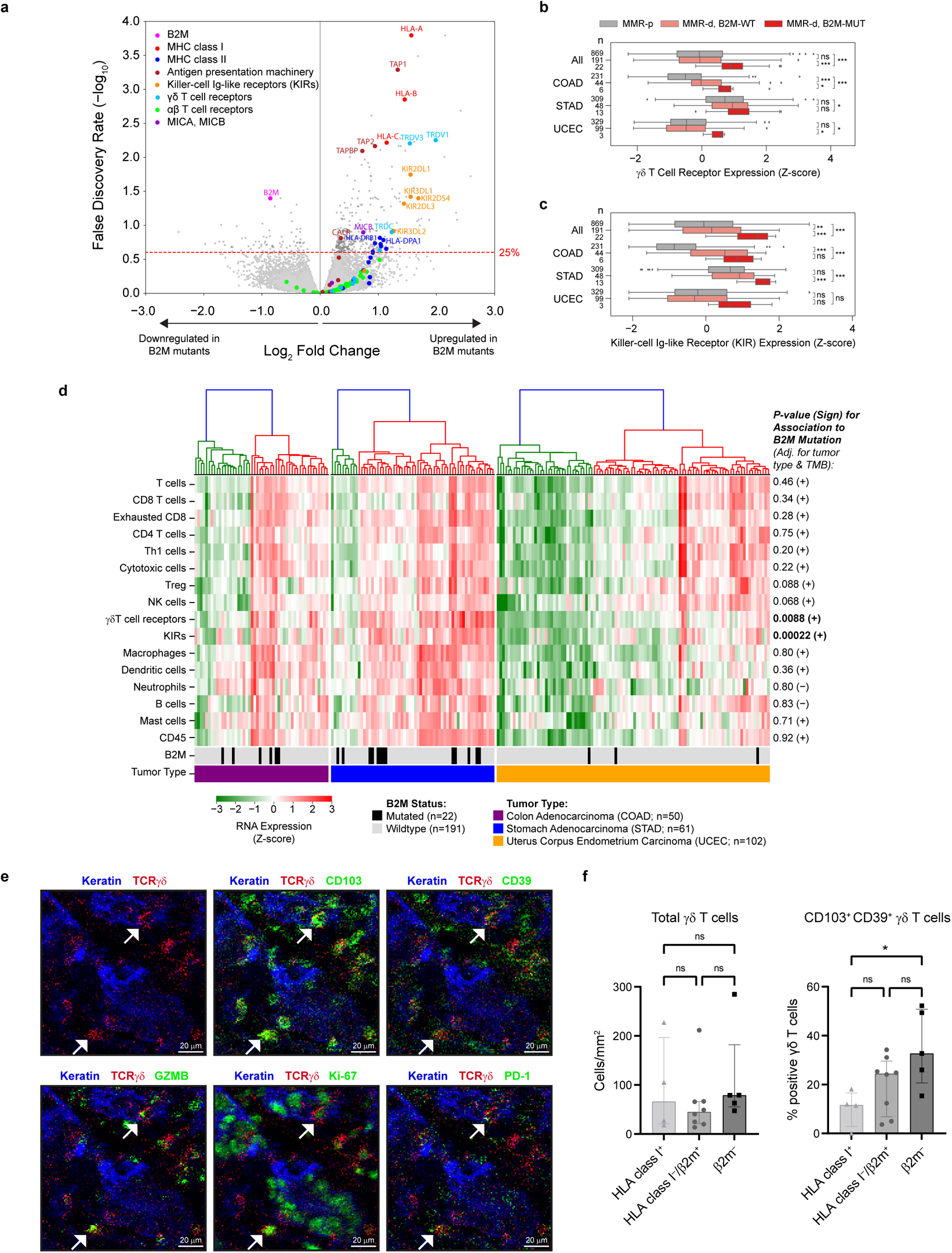
β2m defects are associated with increased infiltration of MMR-d cancers by γδ T cells and killer-cell immunoglobulin-like receptor (KIR)-expressing cells. **a.** Volcano plot indicating differential gene expression between MMR-d cancers with vs. without high impact inactivating mutations in B2M. The Benjamini Hochberg false discovery rate (FDR) significance threshold of 25% is indicated by the red dashed line. Results were obtained in a combined analysis on the TCGA COAD, STAD and UCEC cohorts, and were adjusted for tumor type and tumor mutational burden. **b.** Boxplot showing the RNA expression of γδ T cell receptors in MMR-p (gray), MMR-d B2MWT (pink), and MMR-d B2MMUT (red) cancers. Results are obtained with the TCGA COAD, STAD and UCEC cohorts, and are shown for all cohorts combined (All), and for each cohort separately. Boxes, whiskers, and dots indicate quartiles, 1.5 interquartile ranges, and outliers, respectively. P-values were calculated by Wilcoxon rank sum test. *P<0.05; **P<0.01; ***P<0.001. **c.** As (b), but for RNA expression of killer-cell immunoglobulin-like receptors (KIRs). **d.** Heatmap of the expression (Z-score; see color bar) of gene sets whose expression marks infiltration of specific immune cell types in MMR-d cancers of the COAD, STAD and UCEC cohorts of TCGA. Cancers were ranked based on hierarchical clustering, as indicated by the dendrograms (top). The lower two bars indicate the B2M mutation status and cancer type, as defined in the legend. P-values and sign (+ for positive and – for negative) of associations of the expression of each marker gene set with B2M mutation status are show on the right. P-values were obtained by ordinary least squares linear regression and adjusted for tumor type and tumor mutational burden. Significant associations are in bold font. **e.** Representative images of the detection of tissue-resident (CD103+), activated (CD39+), cytotoxic (granzyme B+), proliferating (Ki-67+), and PD-1+ γδ T cells by imaging mass cytometry in an ICB-naïve, MMR-d colon cancer with β2m defect. **f.** Frequencies of total γδ T cells and CD103+ CD39+ γδ T cells in ICB-naïve HLA class I-positive (+) (n=4), HLA class I-negative (–)/β2m+ (n=8), and β2m– MMR-d colon cancers (n=5). Bars indicate median ± IQR. Each dot represents an individual sample. P-values were calculated by Kruskal-Wallis test with Dunn’s test for multiple comparisons. *P<0.05.

We next assessed if the upregulation of γδ TCRs and KIRs in *B2M^MUT^* MMR-d tumors was caused by higher overall levels of cellular infiltration. To this end, we used marker gene sets^19^ to estimate the abundance of a broad set of immune cell types based on the RNA expression data of the TCGA cohorts. Hierarchical clustering identified a highly and a lowly infiltrated cluster in each of the three tumor types (Fig. 1d). Apart from the γδ TCR and KIR gene sets, none of the other marker gene sets showed increased expression in *B2M^MUT^* versus *B2M^WT^* tumors (Fig. 1d).

Imaging mass cytometry analysis of MMR-d colon cancers, with HLA class I-loss due to defects in β2m, revealed that γδ T cells frequently displayed an intraepithelial localization and expression of CD103 (tissue-residency), CD39 (activation), granzyme B (cytotoxicity), and Ki-67 (proliferation), as well as PD-1 (Fig. 1e). Interestingly, β2m^−^ cancers showed a significantly increased fraction of CD103^+^CD39^+^ γδ T cells as compared to HLA class I^+^ cancers (two-sided P=0.0307 by Kruskal-Wallis test; Fig. 1f). Co-expression of CD103 and CD39 was reported to identify tumor-reactive CD8^+^ αβ T cells in a variety of cancers^20^. Altogether, these data support a role for γδ T cells in mediating natural cytotoxic antitumor responses in HLA class I-negative MMR-d colon cancers.

### Cytotoxic Vδ1 and Vδ3 γδ T cells infiltrate MMR-d colon cancers

To investigate which γδ T cell subsets are present in MMR-d colon cancers and to determine their functional characteristics, we performed single-cell RNA-sequencing on γδ T cells isolated from five MMR-d colon cancers (Extended Data Fig. 2, Extended Data Table 1). Three distinct Vδ subsets were identified (Fig. 2a, Extended Data Fig. 3), where Vδ1 T cells were the most prevalent (43% of γδ T cells), followed by Vδ2 (19%) and Vδ3 T cells (11%) (Fig. 2b). *PDCD1* (encoding PD-1) was predominantly expressed by Vδ1 and Vδ3 γδ T cells, while Vδ1 cells expressed high levels of genes encoding activation markers such as CD39 (*ENTPD1*) and CD38 (Fig. 2c, Extended Data Fig. 2). Furthermore, proliferating γδ T cells (expressing *MKI67*) were especially observed in the Vδ1 and Vδ3 subsets (Fig. 2c). Other distinguishing features of Vδ1 and Vδ3 T cell subsets included the expression of genes encoding activating receptors NKp46 (*NCR1*), NKG2C (*KLRC2*), and NKG2D (*KLRK1*) (Fig. 2c). Interestingly, the expression of several KIRs was also higher in the Vδ1 and Vδ3 subsets as compared to Vδ2 T cells (Fig. 2c). Almost all γδ T cells displayed expression of genes encoding Granzyme B (*GZMB*), Perforin (*PRF1*), and Granulysin (*GNLY*) (Fig. 2c).

**Figure 2.**
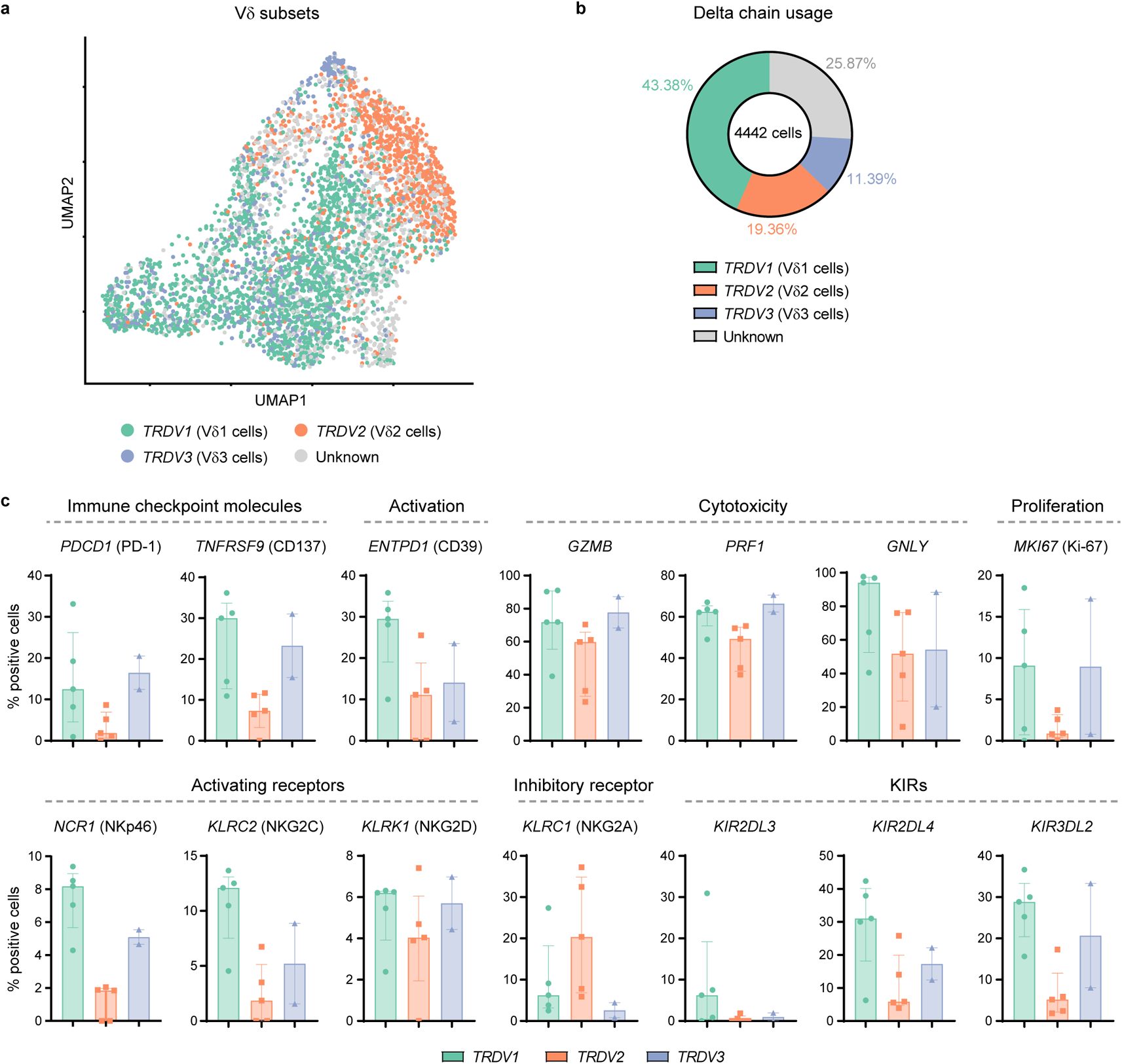
Tumor-infiltrating Vδ1 and Vδ3 T cell subsets display hallmarks of cytotoxic activity in MMR-d colon cancers. **a.** UMAP embedding showing the clustering of γδ T cells (n=4442) isolated from MMR-d colon cancers (n=5) analyzed by single-cell RNA-sequencing. Colors represent the TCR Vδ chain usage. The functionally distinct γδ T cell clusters are shown in Extended Data Fig. 3. Each dot represents a single cell. **b.** Frequencies of the TCR Vδ chain usage of the γδ T cells (n=4442) analyzed by single-cell RNA-sequencing as a percentage of total γδ T cells. **c.** Frequencies of positive cells for selected genes across Vδ1 (n=1927), Vδ2 (n=860), and Vδ3 (n=506) cells as percentage of total γδ T cells from each MMR-d colon tumor (n=5) analyzed by single-cell RNA-sequencing. Vδ3 cells were present in two out of five colon cancers. Bars indicate median ± IQR. Each dot represents an individual sample.

**Figure 3.**
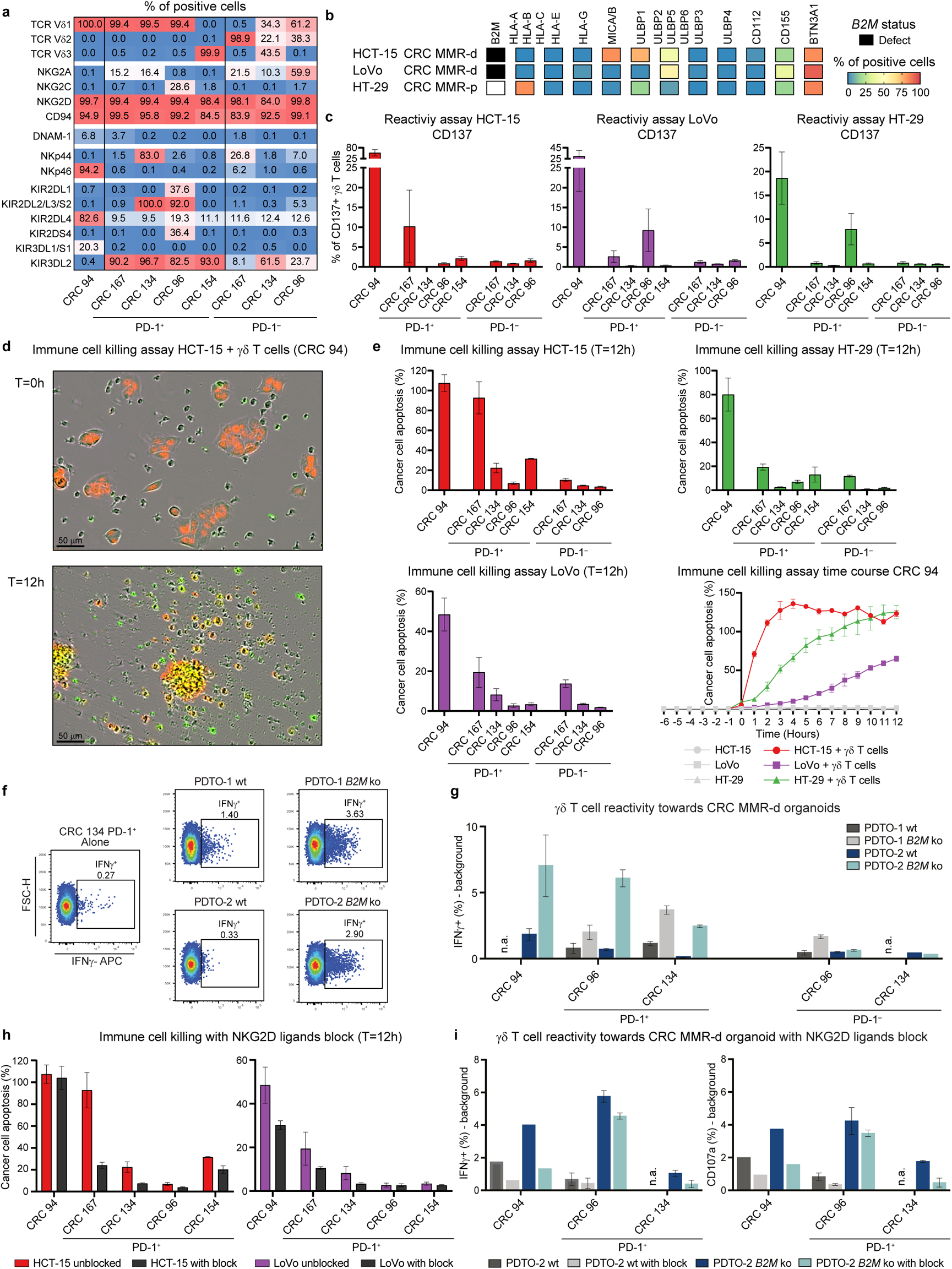
γδ T cells from MMR-d colon cancers show preferential reactivity towards HLA class I-negative cancer cell lines and organoids, which is regulated by NKG2D/NKG2D ligand interactions. **a.** Table showing the percentage of positive cells for different TCR Vδ chains, innate immune receptors, and KIRs on expanded PD-1+ and PD-1– γδ T cells sorted from MMR-d colon cancers (n=5) as percentage of total γδ T cells. **b.** Diagram showing the B2M mutational status and surface expression of HLA class I, NKG2D ligands, DNAM-1 ligands, and butyrophilin on CRC cell lines HCT-15, LoVo, and HT-29. **c.** Bar plots showing the percentage of CD137-positive γδ T cells after 18h co-culture of PD-1+ and PD-1– γδ T cells from MMR-d colon cancers (n=5) with HCT-15, LoVo, and HT-29 cells. Medium as negative control and PMA/ionomycin as positive control are shown in Extended Data Fig. 5. Bars indicate mean ± SEM. Data from four (CRC94), three (CRC167, CRC96), or two (CRC134, CRC154) independent experi­ments, depending on availability of γδ T cells. **d.** Representative images showing the killing of NucLight Red-transduced HCT-15 cells by γδ T cells (unla­beled) from CRC94 in the presence of a green fluorescent caspase-3/7 reagent in the IncuCyte S3. Images are taken immediately after the addition of γδ T cells (T=0) and 12h after. Cancer cell apoptosis is visualized in yellow. **e.** Bar plots showing the quantification of the killing of cancer cell lines by γδ T cells from MMR-d colon cancers (n=5) as in (d) after 12h co-culture. Bars indicate mean ± SEM of two wells with two images/well. At lower right, representative time course of cancer cell apoptosis in the presence or absence of γδ T cells derived from CRC94. **f.** Representative flow cytometry plots of PD-1+ γδ T cells from CRC134 indicating IFNγ expression in unstimulated condition (alone) and upon stimulation with two B2MWT and B2MKO CRC MMR-d organoids, as specified in the subplot titles. **g.** Histogram showing IFNγ expression of γδ T cells from MMR-d colon cancers upon stimulation with two B2MWT and B2MKO CRC MMR-d organoids, as specified in the legend. Background IFNγ signal of each unstimulated γδ T cell sample was subtracted from tumor organoid-stimulated IFNγ signal. For all γδ T cell samples, data is shown for two biological replicates except for CRC134 PD-1– (n=1). Whiskers indicate SEM. **h.** Bar plots showing the quantification of the killing of cancer cell lines by γδ T cells from MMR-d colon cancers (n=5) in the presence of blocking antibodies for NKG2D ligands as compared to the unblocked condition after 12h co-culture. Bars indicate mean ± SEM of two wells with two images/well. **i.** Histograms showing IFNγ (left) and CD107a (right) expression in γδ T cells from MMR-d colon cancers upon stimulation with B2MWT PDTO-2 (gray shades) or B2MKO PDTO-2 (blue shades), with or without NKG2D ligand blocking (as indicated in the legend). For cultured γδ T cells, data is shown for two biological replicates (n=2) except for CRC94 (n=1). Whiskers indicate SEM.

### PD-1^+^ γδ T cells are cytotoxic towards HLA class I-negative colon cancer cells

We next sought to determine whether tumor-infiltrating γδ T cells can recognize and kill CRC cells. We isolated and expanded PD-1^−^ and PD-1^+^ γδ T cells (Extended Data Fig. 4) from five MMR-d colon cancers (Extended Data Table 1). In line with the scRNA-seq data, expanded PD-1^+^ γδ T cells were devoid of Vδ2^+^ cells and comprised of Vδ1^+^ or Vδ3^+^ subsets, whereas PD-1^−^ fractions contained Vδ2^+^ or a mixture of Vδ1/Vδ2/Vδ3^+^ populations (Fig. 3a, Extended Data Fig. 4). Detailed immunophenotyping of the expanded γδ T cells showed that all cells expressed the activating receptor NKG2D, while the surface expression of KIRs was most frequent on PD-1^+^ γδ T cells (Vδ1 or Vδ3^+^), in line with the scRNA-seq results of unexpanded populations (Fig. 3a, Extended Data Fig. 5).

**Figure 4.**
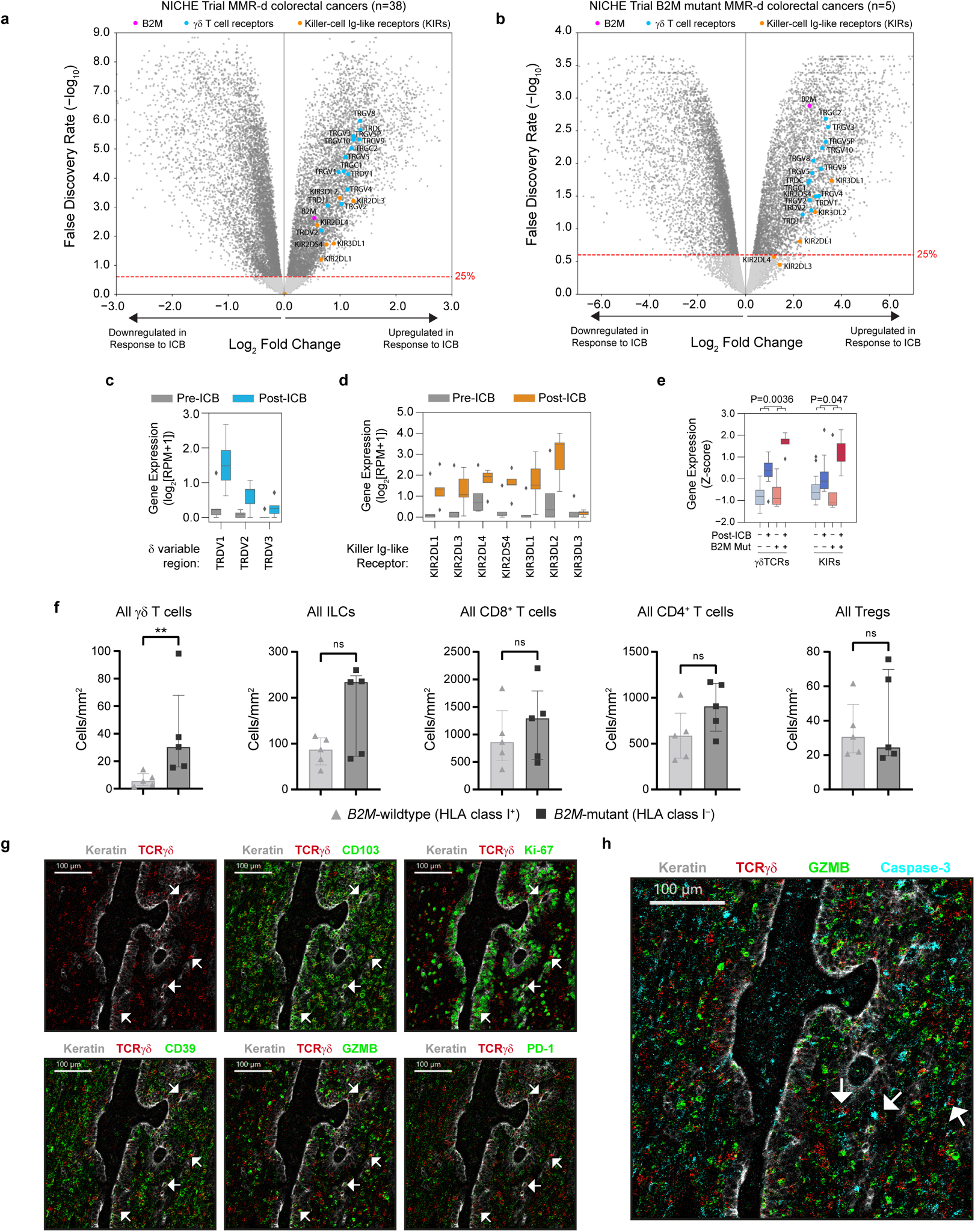
Immune checkpoint blockade (ICB) induces profound infiltration of γδ T cells into MMR-d colon cancers with antigen presentation defects. **a.** Volcano plot indicating differential RNA expression of genes between MMR-d cancers before and after ICB in the NICHE study. The Benjamini Hochberg FDR significance threshold of 25% is indicated by the red dashed line. **b.** As (a), but restricted to the five MMR-d cancers in the NICHE study with high impact (inactivating) mutations in B2M. **c.** Boxplot showing the pre- (gray) and post-ICB (blue) RNA expression of δ T cell receptor variable regions in MMR-d cancers in the NICHE study with high impact (inactivating) mutations in B2M. Boxes, whiskers, and dots indicate quartiles, 1.5 interquartile ranges, and outliers, respectively. P-values were calculated by Wilcoxon rank sum test. **d.** As (c), but for killer-cell Ig-like receptors (KIRs; post-ICB in orange). **e.** Boxplot showing the pre- and post-ICB RNA expression of γδ T cell receptors and killer-cell Ig-like receptors for MMR-d cancers with and without high impact B2M mutations (as indicated in below the x-axis). P-values were for B2M_status:treatment interaction in an ordinary least squares linear regression model. Significant p-values indicate that the treatment-induced increase in γδ TCR/KIR expression is more pronounced in B2MMUT versus B2MWT cancers. Boxes, whiskers, and dots indicate quartiles, 1.5 interquartile ranges, and outliers, respectively. **f.** Frequencies of different immune cell populations in B2MWT (HLA class I-positive, n=5) and B2MMUT (HLA class I-negative, n=5) MMR-d colon cancers upon ICB treatment. Bars indicate median ± IQR. Each dot represents an individual sample. P-values were calculated by Mann-Whitney test. **P<0.01 g. Representative images of the detection of tissue-resident (CD103+), proliferating (Ki-67+), activated (CD39+), cytotoxic (GZMB+), and PD1+ γδ T cells by imaging mass cytometry in a B2MMUT MMR-d colon cancer upon ICB treatment. **h.** Representative image showing the interaction between γδ T cells (colored in red) and caspase-3+ cancer cells (colored in cyan) by imaging mass cytometry in a B2MMUT MMR-d colon tumor upon ICB treatment.

We measured the reactivity of the expanded γδ T cell populations towards HLA class I-negative and HLA class I-positive cancer cell lines (Fig. 3b, Extended Data Fig. 5). Upon co­culture with the different cancer cell lines, expression of activation markers and secretion of IFNγ was mainly induced in PD-1^+^ γδ T cells (Vδ1 or Vδ3^+^) and cell reactivity was most pronounced against HLA class I-negative cell lines (Fig. 3c, Extended Data Fig. 5–6). Reactivity of PD-1^−^ (enriched in Vδ2^+^) subsets towards colorectal cancer cell lines was not detected (Fig. 3c, Extended Data Fig. 5–6). To quantify and visualize the differences in killing of CRC cell lines by PD-1^+^ and PD-1^−^ γδ T cells, we co-cultured the γδ T cell populations with three CRC cell lines (HCT-15, LoVo, HT-29) transduced with a fluorescent caspase-3/7 reagent to measure cancer cell apoptosis over time (Fig. 3d-e). This showed pronounced cancer cell apoptosis upon co-culture with PD-1^+^ γδ T cells (Vδ1 or Vδ3^+^) compared to PD-1^−^ cells, with highest killing of HLA class I-negative HCT-15 cells (Fig. 3e, Movie 1-2).

Next, we established two parental patient-derived tumor organoid lines (PDTOs; Extended Data Table 2) of MMR-d CRC and generated isogenic *B2M^KO^* lines using CRISPR. Genomic knockout of *B2M* effectively abrogated cell surface expression of HLA class I(Extended Data Fig. 7). We exposed two *B2M^KO^* and their parental *B2M^WT^* lines to the expanded γδ T cell subsets, and quantified γδ T cell activation by determination of IFNγ expression. Similarly to our cell line data, γδ T cells displayed increased reactivity towards *B2M^KO^* PDTOs in comparison to the *B2M^WT^* PDTOs (Fig. 3f-g). Furthermore, γδ T cell reactivity towards *B2M^KO^* tumor organoids was preferentially contained within the PD-1^+^ population of γδ T cells (Fig. 3g). Thus, lack of HLA class I antigen presentation in MMR-d tumor cells can be effectively sensed by γδ T cells and stimulates their antitumor response.

Expression of NKG2D on γδ T cells decreased during co-culture with target cells (Extended Data Fig. 8), indicating the involvement of the NKG2D receptor in γδ T cell activity. The NKG2D ligands MICA/B and ULBPs were expressed by the cancer cell lines (Fig. 3b) and the MMR-d CRC PDTOs, irrespective of their *B2M* status (Extended Data Fig. 7). To explore which receptor-ligand interactions might regulate the activity of PD-1^+^ γδ T cells, we performed blocking experiments focused on (i) NKG2D, (ii) DNAM-1, and (iii) γδ TCR signaling. Of these candidates, the only consistent inhibitory effect was observed for NKG2D ligand blocking on cancer cells, which decreased the activation and killing capacity of most PD-1^+^ γδ T cells (Fig. 3h, Extended Data Fig. 9–10), confirming the mechanistic involvement of the NKG2D receptor in γδ T cell activation in this context. In addition, blocking NKG2D ligands on MMR-d CRC PDTOs reduced the PDTO-directed tumor reactivity of γδ T cells from CRC94 and CRC134 (Fig. 3i). Together, these results show that γδ T cell reactivity towards MMR-d tumors is partly dependent on NKG2D/NKG2D-ligand interactions.

### Activated **γ**δ T cells infiltrate ICB-treated *B2M*-mutant MMR-d colon cancers

Next, we studied how ICB influences γδ T cell infiltration and activation in MMR-d colon cancers in a therapeutic context. For this purpose, we analysed pre-and post-treatment samples of the NICHE trial^11^, in which colon cancer patients were treated with neoadjuvant PD-1 plus CTLA-4 blockade. In MMR-d colon cancers (n=38), genes encoding γδ TCRs were highly upregulated in response to ICB (Fig. 4a). When specifically focusing on the five *B2M^MUT^* cancers, we found that expression of genes encoding γδ TCRs was also strongly induced upon ICB, with consistently larger effect sizes as compared to *B2M^WT^* cancers (Fig. 4b). Of the δ variable regions, expression of *TRDV1* (Vδ1) was most strongly induced upon ICB in *B2M^MUT^* cancers (Fig. 4c). The expression of KIRs was also upregulated upon treatment with ICB, both in the cohort as a whole, and in the subgroup of *B2M^MUT^* cancers (Fig. 4a-b). The set of KIRs upregulated upon ICB in *B2M^MUT^* cancers (Fig. 4d) was consistent with the sets of KIRs upregulated in *B2M^MUT^* MMR-d cancers in TCGA (Fig. 1a), and those expressed by MMR-d tumor-infiltrating γδ T cells (Fig. 2c). Finally, expression of the γδ TCR and KIR gene sets was more strongly induced upon response to ICB in *B2M^MUT^* versus *B2M^WT^* cancers (γδ TCRs: two-sided interaction P=0.0056; KIRs: two-sided interaction P=0.047 Fig. 4e).

To quantify and investigate differences in phenotype of γδ T cells upon ICB treatment, we applied imaging mass cytometry to profile immune cell infiltration in post-ICB tissues derived from five *B2M^MUT^* HLA class I-negative and five *B2M^WT^* HLA class I-positive cancers. Due to major pathologic clinical responses, residual cancer cells were absent in most post-ICB treatment samples. All tissues showed a profound infiltration of different types of immune cells, of which γδ T cell infiltration was significantly increased in ICB-treated *B2M^MUT^* as compared to *B2M^WT^* MMR-d colon cancers (two-sided P=0.0079 by Mann-Whitney test; Fig. 4f, Extended Data Fig. 11). In the sole *B2M^MUT^* case that contained cancer cells, γδ T cells displayed co-expression of CD103, Ki-67, CD39, granzyme B, and PD-1 (Fig. 4g). Furthermore, we observed that γδ T cells directly interacted with caspase-3^+^ apoptotic cancer cells in this tumor (Fig. 4h). Taken together, these results show that ICB treatment of MMR-d colon cancer profoundly increases the intratumoral presence of activated, cytotoxic and proliferating γδ T cells, especially when these cancers are β2m-deficient, pointing out γδ T cells as effectors of ICB treatment within this context.

## Discussion

CD8^+^ αβ T cells are major effectors of ICB^13,14,21^, but the clinically relevant responses observed in MMR-d cancers with HLA class I-defects suggest the involvement of other immune effector cells. Here, we show that genomic inactivation of *B2M* in MMR-d colon cancers was associated with: (i) an elevated frequency of activated γδ T cells in ICB-naïve tumors, (ii) an increased presence of tumor-infiltrating γδ T cells upon ICB treatment, (iii) *in vitro* activation of tumor-infiltrating γδ T cells by colorectal cancer cell lines and PDTO, and iv) killing of these tumor cells by γδ T cells, in particular by Vδ1 and Vδ3 subsets expressing PD-1.

Different subsets of γδ T cells exhibit remarkably diverse functions which, in the context of cancer, ranges from tumor-promoting to tumoricidal effects.^22,23^ Hence, it is of interest what defines antitumor reactivity of γδ T cells. Our data suggest that especially tumor-infiltrating Vδ1 and Vδ3 T cells can recognize and kill HLA class I-negative MMR-d tumors, whereas Vγ9Vδ2 cells, the most studied and main subset of γδ T cells in the blood, appear to be less relevant within this context. This is in line with other studies showing that the cytotoxic ability of Vδ1 cells generally outperforms their Vδ2 counterparts^24–28^. Of note, cytotoxicity of tumor-infiltrating Vδ3 cells has, to our knowledge, not been reported before. Furthermore, the observation that PD-1^+^ γδ T cells demonstrated clearly higher levels of antitumor reactivity as compared to their PD-1^−^ counterparts suggests that, as for CD8^+^ αβ T cells^29^, PD-1 expression may be a marker of antitumor reactivity in γδ T cells.

The mechanisms of activation of γδ T cells are notoriously complex and diverse^16^. Specifically for Vδ1^+^ cells, NKG2D has been described to be involved in tumor recognition, which is dependent on tumor cell expression of NKG2D ligands MICA/B and ULBPs^30,31^. In our study, MICA/B and ULBPs were highly expressed by the MMR-d CRC cell lines and tumor organoids, and blocking these ligands reduced γδ T cell activation and cytotoxicity. This suggests a role for the activation receptor NKG2D in γδ T cell reactivity towards HLA class I-negative MMR-d tumors. In addition, we detected expression of KIRs primarily on PD-1^+^ γδ T cells (Vδ1 or Vδ3^+^ subsets), whose antitumor reactivity and killing was clearly amplified when tumor cells lacked HLA class I.

Our findings have broad implications for cancer immunotherapy. First, our results suggest that MMR-d cancers and other tumors with HLA class I defects may be particularly attractive targets for Vδ1 or Vδ3 γδ T cell-based cellular therapies. Second, our findings provide a basis for novel (combinatorial) immunotherapeutic approaches to further enhance γδ T cell-based antitumor immunity. Third, the presence or absence in tumors of specific γδ T cell subsets (e.g. Vδ1 or Vδ3) may help to define patients (un)responsive to ICB, especially in the case of MMR-d cancers and other malignancies with frequent HLA class I defects, like stomach adenocarcinoma^32^ and Hodgkin lymphoma^33^.

Although we have provided detailed and multidimensional analyses, our study is relatively small and it is conceivable that γδ T cells are not the only factor driving ICB responses in HLA class I-negative MMR-d CRC tumors. In this context, other HLA class I-independent immune subsets, like innate lymphoid cells (ILCs), (neoantigen-specific) CD4^+^ T cells (as reported in murine MMR-d cancer models^34^), and macrophages may also contribute. In addition, the requirements of other immune cell types in providing help for effector functions of γδ T cells needs to be clarified.

In conclusion, our results provide strong evidence that γδ T cells are cytotoxic effector cells of ICB treatment in HLA class I-negative MMR-d colon cancers, with implications for further exploitation of γδ T cells in cancer immunotherapy.

## Methods

### TCGA data

RNA expression data (raw counts and Fragments Per Kilobase of transcript per Million mapped reads upper quartile FPKM-UQ) of the colon adenocarcinoma (COAD), stomach adenocarcinoma (STAD) and Uterus Corpus Endometrium Carcinoma (UCEC) cohorts of The Cancer Genome Atlas (TCGA) Research Network were downloaded via the GDC data portal (https://portal.gdc.cancer.gov) on April10^th^, 2019. Of these cohorts, mutation calls of TCGA’s final project, the PanCanAtlas, were downloaded from Synapse (syn7824274) on September 18^th^, 2017. These mutation calls were generated in a standardized pipeline across all samples, resulting in a uniform dataset. Mismatch repair-deficiency status was obtained from Thorsson *et al*^35^ (TCGA Subtype = GI.HM-indel or UCEC.MSI).

### NICHE study sequencing data

Raw RNA reads (FASTA files) of our recently published NICHE study^10^ (ClinicalTrials.gov: <u>NCT03026140</u>) were generated as described in the original publication and aligned to the human reference genome (GRCh38) with STAR software^36^, version 2.7.7a, using default settings. For gene expression quantification, we used the gencode.v35.annotation.gtf annotation file. Somatic mutation data were obtained from DNA sequencing of pre-treatment tumor biopsies and matched germline DNA, as described in the original publication^10^.

### Differential expression analysis

Differential RNA expression of genes was tested in R using EdgeR^37^, Limma^38^ and Voom^39^. Raw read counts were filtered by removing lowly expressed genes. Normalization factors were calculated using EdgeR, in order to transform the raw counts to log_2_ counts per million reads (CPM) and calculate residuals using Voom. Voom was then used to fit a smoothened curve to the √(residual standard deviation) by average gene expression, which was then plotted for visual inspection to confirm that the appropriate threshold was used for filtering of lowly expressed genes (defined as the minimal amount of filtering necessary to overcome a dipping mean-variance trend at low counts). Next, Limma was used to calculate differential expression of genes based on a linear model fit, considering the smoothened curve for sample weights, and empirical Bayes smoothing of standard errors. False discovery rates (FDRs) were calculated by Benjamini-Hochberg correction of the obtained p-values.

#### TCGA data

Using TCGA data, we calculated differential expression between tumors with and without high impact mutations in B2M, adjusting for tumor type and tumor mutational burden (TMB), using the following design formula: expression ∼ Primary_Site + TMB + B2M_status (+ intercept by default), for which Primary_Site was a three-leveled factor (COAD, STAD, or UCEC), TMB was a continuous variable (log_10_[exome-wide number of mutations]) and B2M_status was a two-leveled factor (mutated, or wildtype).

#### NICHE study data

Using NICHE study data, we calculated differential expression between pre- and post-ICB treatment. In order to respect the paired nature of these data, we used the following design formula: expression ∼ Patient + ICB (+ intercept by default), for which Patient was a factor for each individual patient and ICB was a two-leveled factor (ICB-treated yes/no).

### Immune marker gene set expression analysis

#### TCGA data

To utilize RNA-seq data in order to obtain a relative estimate of the infiltration of specific immune cell types within tumors of TCGA, we summed the log_2_(FPKM-UQ+1) expression of genes that are specifically expressed in the immune cell types of interest. To this end, we used the marker gene sets published by Danaher et al.^19^, and extended this by (i) the CD4 T cell marker genes of Davoli et al.^40^, (ii) a γδT cell receptor gene set (comprised of all genes whose name starts with “TRDC”, “TRGC”, “TRDV”, “TRGV”, “TRDJ”, “TRGJ”), and (iii) a killer-cell Ig­like receptor (KIR) gene set (comprised of all genes whose name starts with “KIR” and whose name contains “DL” or “DS”. We excluded the gene set “NK CD56dim cells” of Danaher et al. (comprising IL21R, KIR2DL3, KIR3DL1, and KIR3DL2) from our analyses, as three out of four genes within this set were KIRs and hence this set showed high collinearity/redundancy to our full KIR gene set. The gene set-specific expression values were Z-score transformed. For TCGA-based analyses of MMR-d tumors, association of marker gene set expression with B2M mutation status (high impact mutation yes/no) was calculated using ordinary least squares linear regression, as implemented in the Python package Statsmodels (https://pypi.org/project/statsmodels/), adjusting for tumor type and TMB as described above for the TCGA differential expression analysis.

#### NICHE study data

Based on NICHE study data, we tested if expression of the γδT cell receptor gene set and killer-cell Ig-like receptor (KIR) gene set were more strongly induced upon ICB-treatment in *B2M^MUT^* MMR-d tumors as compared to *B2M^WT^* MMR-d tumors. First, we summed the log_2_(reads per million+1) expression of the genes within the two concerning gene sets for each sample. Next, we fitted an ordinary least squares linear regression model (Statsmodels, see above) that respects the paired nature of the data, using the following design formula: expression ∼ Patient + ICB + B2M + ICB:B2M + intercept, were Patient was a factor for each individual patient, ICB was a two-leveled factor (ICB-treated yes/no), *B2M* was a two-leveled factor (*B2M^MUT^*/*B2M^WT^*), and ICB:B2M was an interaction term between ICB and *B2M*, which represents the statistic of interest (is ICB-based induction of expression of the two gene sets significantly different between *B2M^MUT^* and *B2M^WT^* patients).

### Hierarchical clustering

Hierarchical clustering of immune marker gene set expression profiles (Z-scores) of TCGA cohorts was performed using the Python package Scipy^41^, with Euclidean distance as distance metric and using the Ward variance minimization algorithm.

### Patient samples

Primary colon cancer tissues were from 17 patients with colon cancer who underwent surgical resection of their tumor at the Leiden University Medical Center (LUMC, the Netherlands) (Extended Data Table 1). No patient with a previous history of inflammatory bowel disease was included. This study was approved by the Medical Ethical Committee of the Leiden University Medical Center (protocol P15.282), and patients provided written informed consent. In addition, primary colon cancer tissues from 10 patients with colon cancer included in the NICHE study (<u>NCT03026140</u>)^10^ carried out at the Netherlands Cancer Institute (NKI, the Netherlands) were used for this study. All specimens were anonymized and handled according to the ethical guidelines described in the Code for Proper Secondary Use of Human Tissue in the Netherlands of the Dutch Federation of Medical Scientific Societies.

### Processing of colorectal cancer tissues

Details on the processing of colorectal tumor tissues have been described previously^18^. In short, macroscopic sectioning from the lumen to the most invasive area of the tumor was performed. Tissues were collected in IMDM+Glutamax medium (Gibco) complemented with 20% fetal calf serum (FCS) (Sigma-Aldrich), 1% pen/strep (Gibco) and fungizone (Gibco), and 0.1% ciprofloxacin (provided by apothecary LUMC) and gentamicin (Invitrogen), and immediately cut into small fragments in a petri dish. Enzymatical digestion was performed with 1 mg/mL collagenase D (Roche Diagnostics) and 50 µg/mL DNase I (Roche Diagnostics) in 5 mL of IMDM+Glutamax medium for 30 min at 37°C in gentleMACS C tubes (Miltenyi Biotec). During and after incubation, cell suspensions were dissociated mechanically on the gentleMACS Dissociator (Miltenyi Biotec). Cell suspensions were filtered through a 70-µm cell strainer (Corning), washed in IMDM+Glutamax medium with 20% FCS, 1% pen/strep, and 0.1% fungizone, and cell count and viability were determined with the Muse Count & Viability Kit (Merck) on the Muse Cell Analyser (Merck). Based on the number of viable cells, cells in IMDM+Glutamax medium were cryopreserved in liquid nitrogen until time of analysis complemented 1:1 with 80% FCS and 20% dimethyl sulfoxide (DMSO) (Merck).

### Immunohistochemical detection of MMR, β2m, and HLA class I proteins

Tumor MMR status was determined by immunohistochemical detection of PMS2 (anti-PMS2 antibodies; clone EP51, DAKO) and MSH6 (anti-MSH6 antibodies; clone EPR3945, Abcam) proteins^42^. MMR-deficiency was defined as the lack of expression of at least one of the MMR-proteins in the presence of an internal positive control. Tumor β2m status was determined by immunohistochemical detection of β2m (anti-β2m antibodies; clone EP2978Y, Abcam). Immunohistochemical detection of HLA class I expression on tumors was performed with HCA2 and HC10 monoclonal antibodies (Nordic-MUbio), and classified as HLA class I positive, weak, or loss as described previously^6^.

### Imaging mass cytometry staining and analysis

Imaging mass cytometry (IMC) was performed on ICB-naïve colon cancer tissues (MMR-d) of 17 patients from the LUMC, of which four HLA class I-positive, eight HLA class I-defect, and five β2m-defect (Extended Data Table 1). In addition, IMC was performed on ICB-treated colon cancer tissues (MMR-d) of ten patients from the NKI, of which five *B2M^WT^* and five *B2M^MUT^*. Antibody conjugation and immunodetection were performed following the methodology published previously by Ijsselsteijn *et al.*^43^. Four-µm FFPE tissue were incubated with 41 antibodies in four steps. First, sections were incubated with anti-CD4 and anti-TCRδ overnight at RT, which were subsequently detected using metal-conjugated secondary antibodies (goat anti-rabbit IgG and goat anti-mouse IgG, respectively; Abcam). Second, sections were incubated with 20 antibodies (Extended Data Table 3) for five hours at RT. Third, sections were incubated overnight at 4°C with the remaining 19 antibodies (Extended Data Table 3). Fourth, sections were incubated with 0.125 µM Cell-ID intercalator-Ir (Fluidigm) to detect the DNA, and stored dry until measurement. For each sample, six 1000×1000µm regions were selected based on consecutive Haematoxylin and Eosin (H&E) stains and ablated using the Hyperion Imaging system (Fluidigm). Data was acquired with the CyTOF Software (version 7.0) and exported with MCD Viewer (version 1.0.5). Data was normalised using semi-automated background removal in ilastik^44^, version 1.3.3, to control for variations in signal-to-noise between FFPE sections as described previously^45^. Thereafter, the phenotype data was normalized at pixel level. Cell segmentation masks were created for all CD3- and/or CD7-positive cells in ilastik and CellProfiler^46^, version 2.2.0. In ImaCytE^47^, version 1.1.4, cell segmentation masks and normalized images were combined to generate single-cell FCS files containing the relative frequency of positive pixels for each marker per cell. Cells forming visual neighbourhoods in a t-distributed Stochastic Neighbour Embedding (t-SNE)^48^ embedding in Cytosplore^49^, version 2.3.0, were grouped and exported as separate FCS files. The resulting subsets were imported back into ImaCyte and visualized on the segmentation masks. Expression of immunomodulatory markers was determined as all cells with a relative frequency of at least 0.2 positive pixels per cell. Differences in cells/mm^2^ were calculated by Mann-Whitney tests in Graphpad Prism (version 9.0.1).

### Sorting of γδ T cells from colon cancers and single-cell RNA-sequencing

scRNA-seq was performed on sorted γδ T cells from colon cancers (MMR-d) of five patients from the LUMC in the presence of hashtag oligo (HTOs) for sample ID and antibody-derived tags (ADTs) for CD45RA and CD45RO protein expression by CITE-seq^50^. Cells were thawed, rest at 37°C in IMDM (Lonza)/20% FCS for 1h, followed by incubation with human Fc receptor block (BioLegend) for 10 min at 4°C. Thereafter, cells were stained with cell surface antibodies (1:50 anti-CD3-PE [clone SK7, BD Biosciences], 1:160 anti-CD45-PerCP-Cy5.5 [clone 2D1, eBioscience], 1:200 anti-CD7-APC [clone 124-1D1, eBioscience], 1:60 anti-EPCAM-FITC [clone HEA-125, Miltenyi], 1:80 anti-TCRγδ-BV421 [clone 11F2, BD Biosciences], and a 1:1000 near-infrared viability dye [Life Technologies]), 1 µg of TotalSeq-C anti-CD45RA (clone HI100, BioLegend) and 1 µg of anti-CD45RO (clone UCHL1, BioLegend) antibodies, and 0.5 µg of a unique TotalSeq-C CD298/β2M hashtag antibody (clone LNH-94/2M2, BioLegend) for each sample (n=5) for 30 min at 4°C. Cells were washed three times in FACS buffer (PBS (Fresenius Kabi)/1% FCS) and kept cold and dark until cell sorting. Compensation was carried out with CompBeads (BD Biosciences) and ArC reactive beads (Life Technologies). Single, live CD45^+^ EPCAM^−^ CD3^+^ TCRγδ^+^ cells from five colorectal tumors (MMR-d) were sorted on a FACS Aria III 4L (BD Biosciences). After sorting, the samples were pooled.

scRNA-seq libraries were prepared using the Chromium Single Cell 5’ Reagent Kit v1 chemistry (10X Genomics) following the manufacturer’s instructions. The construction of 5’ Gene Expression libraries allowed the identification of γδ T cell subsets according to Vδ and Vγ usage. Libraries were sequenced on a HiSeq X Ten using paired-end 2×150 bp sequencing (Illumina). Reads were aligned to the human reference genome (GRCh38) and quantified using Cell Ranger (version 3.1.0). Downstream analysis was performed using Seurat (version 3.1.5) according to the author’s instructions^51^. Briefly, cells that had less than 200 detected genes and genes that were expressed in less than six cells were excluded. The resulting 5669 cells were demultiplexed based on HTO enrichment using the MULTIseqDemux algorithm^52^.

Next, cells with a mitochondrial gene content greater than 10% and cells with outlying numbers of expressed genes (>3000) were filtered out from the analysis, resulting in a final dataset of 4442 cells. Data were normalized using the ‘LogNormalize’ function from Seurat with scale factor 10,000. Variable features were identified using the ‘FindVariableFeatures’ function from Seurat returning 2,000 features. We then applied the ‘RunFastMNN’ function from SeuratWrappers split by sample ID to adjust for potential batch-derived effects across samples^53^. Uniform manifold approximation (UMAP)^54^ was used to visualize the cells in a two-dimensional space, followed by the ‘FindNeighbors’ and ‘FindClusters’ functions from Seurat. Data were scaled and heterogeneity associated with mitochondrial contamination was regressed out. Cell clusters were identified by performing differentially expressed gene analysis with the ‘FindAllMarkers’ function with min.pct and logfc.threshold at 0.25. Percentage of *TRDV1* (Vδ1), *TRDV2* (Vδ2), or *TRDV3* (Vδ3) positive cells was determined as the percentage of all cells with an expression level of >1, while <1 for the other TCR Vδ chains. CRC96, 134 and 167 had less than ten *TRDV3*^+^ cells, and were not included in the Vδ3 analysis. Transcripts of Vδ4 *(TRDV4),* Vδ5 *(TRDV5),* and Vδ8 *(TRDV8)* cells were not detected. Percentage of *TRGV1* (Vγ1) – *TRGV11* (Vγ11) positive cells was determined as the percentage of all cells with an expression level of >1, while <1 for the other TCR Vγ chains. Percentage of cells positive for a certain gene was determined as all cells with an expression level of >1.

### Sorting of γδ T cells from colon cancers and cell culturing

γδ T cells from colon cancers (MMR-d) of five patients from the LUMC were sorted for cell culture. Cells were thawed and rest at 37°C in IMDM (Lonza)/10% nHS for 1h. Thereafter, cells were incubated with human Fc receptor block (BioLegend) and stained with cell surface antibodies (1:20 anti-CD3-Am Cyan [clone SK7, BD Biosciences], 1:80 anti-TCRγδ-BV421 [clone 11F2, BD Biosciences], and 1:30 anti-PD-1-PE [clone MIH4, eBioscience] for 45 min at 4°C together with different additional antibodies for immunophenotyping (including 1:10 anti-CD103-FITC [clone Ber-ACT8, BD Biosciences], 1:200 anti-CD38-PE-Cy7 [clone HIT2, eBioscience], 1:60 anti-CD39-APC [clone A1, BioLegend], 1:20 anti-CD45RA-PE-Dazzle594 [clone HI100, Sony], 1:20 anti-CD45RO-PerCP-Cy5.5 [clone UCHL1, Sony], 1:40 anti-TCRαβ­PE-Cy7 [clone IP26, BioLegend], 1:50 anti-TCRVδ1-FITC [clone TS8.2, Invitrogen], or 1:200 anti-TCRVδ2-PerCP-Cy5.5 [clone B6, BioLegend]. A 1:1000 live/dead fixable near-infrared viability dye (Life Technologies) was included in each staining. Cell were washed three times in FACS buffer (PBS/1% FCS) and kept cold and dark until cell sorting. Compensation was carried out with CompBeads (BD Biosciences) and ArC reactive beads (Life Technologies). Single, live CD3^+^ TCRγδ^+^ PD-1^+^ and PD-1^−^ cells from five colorectal tumors (MMR-d) were sorted on a FACS Aria III 4L (BD Biosciences). For CRC94 all γδ T cells were sorted due to the low number of PD-1^+^ cells. γδ T cells were sorted in medium containing feeder cells (1×10^6^/mL), PHA (1 µg/mL; Thermo Fisher Scientific), IL-2 (1000 IU/mL; Novartis), IL-15 (10 ng/mL; R&D Systems), gentamicin (50 µg/mL), and fungizone (0.5 µg/mL). Sorted γδ T cells were expanded in the presence of 1000 IU/mL IL-2 and 10 ng/mL IL-15 for three-four weeks. Purity and phenotype of γδ T cells were assessed by flow cytometry. We obtained a >170,000­fold increase in 3-4 weeks of expansion of γδ T cells (Extended Data Fig. 4e).

### Immunophenotyping of expanded γδ T cells by flow cytometry

Expanded γδ T cells from colon tumors were analyzed by flow cytometry for the expression of TCR Vδ chains, NKG2 receptors, NCRs, KIRs, tissue-residency/activation markers, cytotoxic molecules, immune checkpoint molecules, cytokine receptors, and Fc receptors. Briefly, cells were incubated with human Fc receptor block (BioLegend) and stained with cell surface antibodies (Extended Data Table 4) for 45 min at 4°C, followed by three washing steps in FACS buffer (PBS/1% FCS). Granzyme B and perforin were detected intracellularly using Fixation Buffer and Intracellular Staining Permeabilization Wash Buffer (BioLegend). Compensation was carried out with CompBeads (BD Biosciences) and ArC reactive beads (Life Technologies). Cells were acquired on a FACS LSR Fortessa 4L (BD Biosciences) running FACSDiva software version 9.0 (BD Biosciences). Data were analyzed with FlowJo software version 10.6.1 (Tree Star Inc).

### Cell culture of cancer cell lines

Human colorectal adenocarcinoma cell lines HCT-15 (MMR-d), LoVo (MMR-d), HT-29 (MMR­p), SW403 (MMR-p), and SK-CO-1 (MMR-p) as well as HLA class I deficient human leukemia cell line K-562 and Burkitt lymphoma cell line Daudi were used as targets for reactivity and immune cell killing assays. The cell lines were authenticated by STR profiling and tested for mycoplasma. HCT-15, LoVo, HT-29, K-562, and Daudi cells were maintained in RPMI (Gibco)/10% FCS. SW403 and SK-CO-1 were maintained in DMEM/F12 (Gibco)/10% FCS. All adherent cell lines were trypsinized before passaging.

### Organoid models and culture

Tumor organoids were derived from MMR-d CRC tumor of two patients via resection from the colon, tumor organoid 1, or peritoneal biopsy, tumor organoid 2 (Extended Data Table 2). Establishment of the respective organoid lines from tumor material was performed as previously reported^55,56^. Briefly, tumor tissue was mechanically dissociated and digested with 1.5 mg/mL of collagenase II (Sigma-Aldrich), 10 µg/mL of hyaluronidase type IV (Sigma­Aldrich), and 10 µM Y-27632 (Sigma-Aldrich). Cells were embedded in Cultrex® RGF BME Type 2 (cat no. 3533-005-02, R&D systems) and placed in a 37°C incubator for 20 min. Human CRC organoids medium is composed of Ad-DF+++ (Advanced DMEM/F12 (GIBCO) supplemented with 2 mM Ultraglutamine I (Lonza), 10 mM HEPES (GIBCO), and 100/100 U/mL Penicillin/Streptomycin (GIBCO), 10% Noggin-conditioned medium, 20% R-spondin1-conditioned medium, 1x B27 supplement without vitamin A (GIBCO), 1.25 mM N­acetylcysteine (Sigma-Aldrich), 10 mM nicotinamide (Sigma-Aldrich), 50 ng/mL human recombinant EGF (Peprotech), 500 nM A83-01 (Tocris), 3 µM SB202190 (Cayman Chemicals) and 10 nM prostaglandin E2 (Cayman Chemicals). Organoids were passaged depending on growth every 1–2 weeks by incubating in TrypLE Express (Gibco) for 5–10 min followed by embedding in BME. Organoids were authenticated by SNP array or STR profile and regularly tested for Mycoplasma using Mycoplasma PCR43 and the MycoAlert Mycoplasma Detection Kit (cat no. LT07-318). In the first two weeks of organoid culture, 1x Primocin (Invivogen) was added to prevent microbial contamination. Procedures performed with patient specimens were approved by the Medical Ethical Committee of the Netherlands Cancer Institute – Antoni van Leeuwenhoek hospital (study NL48824.031.14) and written informed consent was obtained from all patients. Mismatch repair status was assessed by standard protocol for the Ventana automated immunostainer for MLH1 clone M1 (Roche), MSH2 clone G219-1129 (Roche), MSH6 clone EP49 (Abcam) and PMS2 clone EP51 (Agilant Technologies). The *B2M^KO^* tumor organoid lines were generated by using sgRNA targeting *B2M* (GGCCGAGATGTCTCGCTCCG), cloned into LentiCRISPR v2 plasmid. The virus was produced by standard method.

### Screening **of cancer cell lines and tumor organoids by flow cytometry**

The cancer cell lines used in the reactivity and killing assays were screened for the expression of HLA class I molecules, NKG2D ligands, DNAM-1 ligands, and butyrophilin by flow cytometry. Briefly, cells were incubated with human Fc receptor block (BioLegend) and stained with the different cell surface antibodies (1:10 anti-CD112-PE [clone R2.525, BD Biosciences], 1:10 anti-CD155-PE [clone 300907, R&D Systems], 1:50 anti-CD277/BTN3A1-PE [clone BT3.1, Miltenyi], 1:100 anti-HLA-A,B,C-FITC [clone W6/32, eBioscience], 1:20 anti-HLA-E­BV421 [clone 3D12, BioLegend], 1:20 anti-HLA-G-APC [clone 87G, BioLegend], 1:300 anti-MICA/B-PE [clone 6D4, BioLegend], 1:10 anti-ULBP1-PE [clone 170818, R&D Systems], 1:20 anti-ULBP2/5/6-PE [clone 165903, R&D Systems], 1:20 anti-ULBP3-PE [clone 166510, R&D Systems], or 1:20 anti-ULBP4-PE [clone 709116, R&D Systems] for 45 min at 4°C. A 1:1000 live/dead fixable near-infrared viability dye (Life Technologies) was included in each staining. Cells were washed three times in FACS buffer (PBS/1% FCS). Compensation was carried out with CompBeads (BD Biosciences) and ArC reactive beads (Life Technologies). Cells were acquired on a FACS Canto II 3L or FACS LSR Fortessa 4L (BD Biosciences) running FACSDiva software version 9.0 (BD Biosciences). Isotype or FMO controls were included to determine the percentage of positive cancer cells. Data were analyzed with FlowJo software version 10.6.1 (Tree Star Inc).

For organoid surface staining, tumor organoids were dissociated into single cells using TrypLE Express (Gibco) washed twice in cold FACS buffer (PBS, 5 mM EDTA, 1% bovine serum antigen) and stained with either 1:20 anti-HLA-A,B,C-PE (clone W6/32, BD Biosciences), 1:100 anti-β2m-FITC (clone 2M2, BioLegend), 1:200 anti-PD-L1 (clone MIH1, eBioscience) and 1:2000 near-infrared (NIR) viability dye (Life Technologies) or isotype controls (FITC, PE or APC) mouse IgG1 kappa (BD Biosciences). For NKG2D ligand expression analysis cells were stained with 1:300 anti-MICA/MICB, 1:10 anti-ULBP1, 1:20 anti-ULBP2/5/6, 1:20 anti-ULBP3, 1:20 anti-ULBP4, and 1:2000 near-infrared (NIR) viability dye (Life Technologies). Tumor cells were incubated for 30 min at 4°C in the dark and washed twice in FACS buffer. All samples were recorded at a Becton Dickinson Fortessa.

### Reactivity assay γδ T cells

Reactivity of γδ T cells to the different cancer cell lines was assessed by a co-culture reactivity assay. γδ T cells were thawed and cultured in IMDM+Glutamax (Gibco)/8% nHS medium with pen (100 IU/mL)/strep (100 µg/mL) in the presence of low-dose IL-2 (25 IU/mL) and IL-15 (5 ng/mL) overnight at 37°C. Cancer cell lines were counted, adjusted to a concentration of 0.5×10^5^ cells/mL in IMDM+Glutamax/10% FCS medium with pen (100 IU/mL)/strep (100 µg/mL), and seeded (100 µL/well) in coated 96-well flat-bottom microplates (Greiner CellStar) (for 5,000 cells/well) overnight at 37°C. The next day, γδ T cells were harvested, counted, and adjusted to a concentration of 1.2×10^6^ cells/mL in IMDM+Glutamax/10% FCS medium. The γδ T cells were added in 50 µL (for 60,000 cells/well) and co-cultured (12:1 E:T ratio) at 37°C for 18h in biological triplicates. Medium (without cancer cells) was used as negative control and PMA (20 ng/mL)/ionomycin (1 µg/mL) as positive control. After co-culture, the supernatant was harvested to detect IFNγ secretion by ELISA (Mabtech) following the manufacturer’s instructions. Additionally, cells were harvested, incubated with human Fc receptor block (BioLegend), and stained with cell surface antibodies (1:100 anti-CD137-APC [clone 4B4-1, BD Biosciences], 1:150 anti-CD226/DNAM-1-BV510 [clone DX11, BD Biosciences], 1:400 anti-CD3-AF700 [clone UCHT1, BD Biosciences], 1:80 anti-CD39-APC [clone A1, BioLegend], 1:10 anti-CD40L-PE [clone TRAP1, BD Biosciences] or 1:30 anti-PD-1-PE [clone MIH4, eBioscience], 1:40 anti-TCRγδ-BV650 [clone 11F2, BD Biosciences], 1:300 anti-NKG2D-PE-Cy7 [clone 1D11, BD Biosciences], and 1:20 anti-OX40-FITC [clone ACT35, BioLegend] for 45 min at 4°C. A 1:1000 live/dead fixable near-infrared viability dye (Life Technologies) was included in each staining. Cells were washed three times in FACS buffer (PBS/1% FCS). Compensation was carried out with CompBeads (BD Biosciences) and ArC reactive beads (Life Technologies). Cells were acquired on a FACS LSR Fortessa X-20 4L (BD Biosciences) running FACSDiva software version 9.0 (BD Biosciences). Data were analyzed with FlowJo software version 10.6.1 (Tree Star Inc). All data are representative of at least two independent experiments.

### Immune cell killing assay γδ T cells

Killing of the different cancer cell lines by γδ T cells was visualized and quantified by a co­culture immune cell killing assay using the IncuCyte S3 Live-Cell Analysis System (Essen Bioscience). HCT-15, LoVo, and HT-29 cells were transduced with IncuCyte NucLight Red Lentivirus Reagent (EF-1α, Puro; Essen BioScience) providing a nuclear-restricted expression of a red (mKate2) fluorescent protein. In short, HCT-15, LoVo and HT-29 were seeded, transduced according to the manufacturer’s instructions, and stable cell populations were generated using puromycin selection. Cancer cell lines were counted, adjusted to a concentration of 1×10^5^ cells/mL in IMDM+Glutamax/10% FCS medium with pen (100 IU/mL)/strep (100 µg/mL), and seeded (100 µL/well) in 96-well flat-bottom clear microplates (Greiner CellStar) (for 10,000 cells/well). The target cell plate was placed in the IncuCyte system at 37°C to monitor for cell confluency for 3 days. On day 2, γδ T cells were thawed and cultured in IMDM+Glutamax/8% nHS medium with pen (100 IU/mL)/strep (100 µg/mL) in the presence of low-dose IL-2 (25 IU/mL) and IL-15 (5 ng/mL) overnight at 37°C. The next day, γδ T cells were harvested, counted, and adjusted to a concentration of 7.2×10^5^ cells/mL in IMDM+Glutamax/10% FCS medium. After aspiration of the medium of the target cell plate, 100 µL of new medium containing 3.75 µM IncuCyte Caspase-3/7 Green Apoptosis Reagent (Essen BioScience) (1.5x final assay concentration of 2.5 µM) was added together with 50 µL of γδ T cells (for 36,000 cells/well). They were co-cultured (4:1 E:T ratio) in the IncuCyte system at 37°C in biological duplicates. Cancer cells alone and cancer cells alone with Caspase-3/7 were used as negative controls. Images (2 images/well) were captured every hour at 20x magnification with the phase, green, and red channels for up to 4 days.

Analysis was performed in the IncuCyte software (version 2020B) for each cancer cell line separately. The following analysis definitions were applied: 1) for HCT-15 cells in the phase channel a minimum area of 200 µm^2^, in the green channel a threshold of 2 GCU, and in the red channel a threshold of 2 RCU, 2) for LoVo and HT-29 cells in the phase channel a minimum area of 200 µm^2^, in the green channel a threshold of 4 GCU, and in the red channel a threshold of 2 RCU. Cancer cell apoptosis was then quantified in the IncuCyte software by counting the total number of Green + Red objects per image normalized (by division) to the total number of Red objects per image after 12h co-culture and displayed as a percentage (mean ± SEM) of two wells with two images/well.

### Tumor organoid recognition assay

For evaluation of tumor reactivity towards *B2M^WT^* and *B2M^KO^* organoids and NKG2D ligand blocking conditions, tumor organoids and γδ T cells were prepared as described previously.^10,55,56^ Two days prior to the experiment organoids were isolated from BME by incubation in 2 mg/mL type II dispase (Sigma-Aldrich) for 15 min before addition of 5 mM ethylendiaminetetraacetic acid (EDTA) and washed with PBS before resuspended in CRC organoid medium with 10 µM Y-27632 (Sigma-Aldrich). Organoids were stimulated with 200 ng/mL IFNγ (Peprotech) 24 hours before the experiment. For the recognition assay and intracellular staining tumor organoids were dissociated into single cells and plated in anti-CD28 (clone CD28.2 eBioscience) coated 96-well U-bottom plates with γδ T cells at a 1:1 target:effector ratio in the presence of 20 µg/mL anti-PD-1 (Merus). As positive control γδ T cells were stimulated with 50 ng/mL of phorbol 12-myristate 13-acetate (Sigma-Aldrich) and 1 µg/mL of ionomycin (Sigma-Aldrich). After 1h of incubation at 37°C, GolgiSTOP (BD Biosciences, 1:1500) and GolgiPlug (BD Biosciences, 1:1000) were added. After 4h of incubation at 37°C, γδ T cells were washed twice in cold FACS buffer (PBS, 5 mM EDTA, 1% bovine serum antigen) and stained with 1:20 anti-CD3-PerCP-Cy5.5 (BD Biosciences), 1:20 anti-TCRγδ-PE (BD Bioscience), 1:20 anti-CD4-FITC (BD Bioscience) (not added in experiments with NKG2D ligand blocking), 1:200 anti-CD8-BV421 (BD Biosciences) and 1:2000 near-infrared (NIR) viability dye (Life Technologies) for 30 min at 4°C. Cells were washed, fixed and stained with 1:40 anti-IFNγ-APC (BD Biosciences) for 30 min at 4°C, using the Cytofix/Cytoperm Kit (BD Biosciences). After two washing steps, cells were resuspended in FACS buffer and recorded at a BD LSRFortessa™ Cell Analyzer SORP flow cytometer with FACSDiVa 8.0.2 (BD Biosciences) software.

### Blocking experiments with cancer cell lines and tumor organoids

Reactivity of and killing by the γδ T cells was examined in the presence of different blocking antibodies to investigate which receptor-ligand interactions are involved. For DNAM-1 blocking, γδ T cells were incubated with 3 µg/mL purified anti-DNAM-1 (clone DX11, BD Biosciences) for 1h at 37°C. For γδ TCR blocking, γδ T cells were incubated with 3 µg/mL purified anti-TCRγδ (clone 5A6.E9, Invitrogen) for 1h at 37°C, of which the clone we used was tested to be best for use in γδ TCR blocking assays^57^. NKG2D ligands were blocked on the cancer cell lines and single cells of tumor organoids by incubating the target cells with 12 µg/mL anti-MICA/B (clone 6D4, BioLegend), 1 µg/mL anti-ULBP1 (clone 170818, R&D Systems), 3 µg/mL anti-ULBP2/5/6 (clone 165903, R&D Systems), and 6 µg/mL anti-ULBP3 (clone 166510, R&D Systems) for 1h at 37°C prior to plating with γδ T cells. After incubation with the blocking antibodies, the γδ T cells were added to cancer cell lines HCT-15, LoVo, and HT-29 as described above with a minimum of two biological replicates per blocking condition. For organoid experiments, 1:50 anti-CD107a-FITC (clone H4A3, BioLegend) was added during incubation.

As a control for Fc-mediated antibody effector functions, γδ T cells alone were incubated with the blocking antibodies in the presence of 2.5 µM IncuCyte Caspase-3/7 Green Apoptosis Reagent (Essen BioScience) in the IncuCyte system at 37°C, and the number of apoptotic γδ T cells was quantified over time.

## Abbreviations

B2M/β2m, β2-microglobulin B2M^KO^, β2-microglobulin-knockout B2M^MUT^, β2-microglobulin-mutant B2M^WT^, β2-microglobulin-wildtype BTN, butyrophilin CML, chronic myelogenous leukemia COAD, colon adenocarcinoma CRC, colorectal cancer ICB, immune checkpoint blockade ICS, intra-cellular staining KIR, killer-cell immunoglobulin-like receptor MMR-d, mismatch repair-deficient MMR-p, mismatch repair-proficient NCR, natural cytotoxicity receptor PDTO, patient-derived tumor organoid scRNA-seq, single-cell RNA-sequencing STAD, stomach adenocarcinoma UCEC, uterus corpus endometrium carcinoma UMAP, uniform manifold approximation and projection

## Index **of supplemental information**

**Extended Data Fig. 1.** Association of *B2M* mutation status with expression of αβ T cell receptors.

**Extended Data Fig. 2.** Characterization of γδ T cells from MMR-d colon cancers by scRNA-seq.

**Extended Data Fig. 3.** Distinct clusters of γδ T cells from MMR-d colon cancers by scRNA-seq.

**Extended Data Fig. 4.** Sorting of PD-1^+^ and PD-1^−^ γδ T cells from MMR-d colon cancers by FACS and their TCR Vδ chain usage.

**Extended Data Fig. 5.** Reactivity of γδ T cells from MMR-d colon cancers towards cancer cell lines.

**Extended Data Fig. 6.** Surface expression of activation markers and secretion of IFNγ upon co-culture of γδ T cells from MMR-d colon cancers with cancer cell lines.

**Extended Data Fig. 7.** Tumor organoid characterization and reactivity assay readout.

**Extended Data Fig. 8.** Surface expression of activating receptor NKG2D by PD-1^+^ γδ T cells from MMR-d colon cancers upon co-culture with cancer cell lines.

**Extended Data Fig. 9.** Killing of cancer cell lines by PD-1^+^ γδ T cells from MMR-d colon cancers in the presence NKG2D ligand, DNAM-1, or γδ TCR blocking.

**Extended Data Fig. 10.** Reactivity towards cancer cell lines by PD-1^+^ γδ T cells from MMR-d colon cancers in the presence of NKG2D ligand blocking.

**Extended Data Fig. 11.** Distribution of immune cell populations in *B2M*-wildtype and *B2M*­mutant colon cancers upon ICB by imaging mass cytometry.

**Extended Data Table 1.** Characteristics of clinical samples from 17 patients with MMR-d colon cancer.

**Extended Data Table 2.** Characteristics of patient-derived organoids from MMR-d colorectal cancer.

**Extended Data Table 3.** Antibodies used for imaging mass cytometry of colon cancers.

**Extended Data Table 4.** Antibodies used for immunophenotyping of γδ T cells by flow cytometry.

**Movie 1.** Killing of HCT-15 cells by γδ T cells (Vδ1^+^) from a MMR-d colon cancer.

**Movie 2.** Killing of HCT-15 cells by PD-1^+^ (Vδ1^+^) as compared to PD-1^−^ (Vδ2^+^) γδ T cells from a MMR-d colon cancer.

---

**Extended Data Fig. 1.**
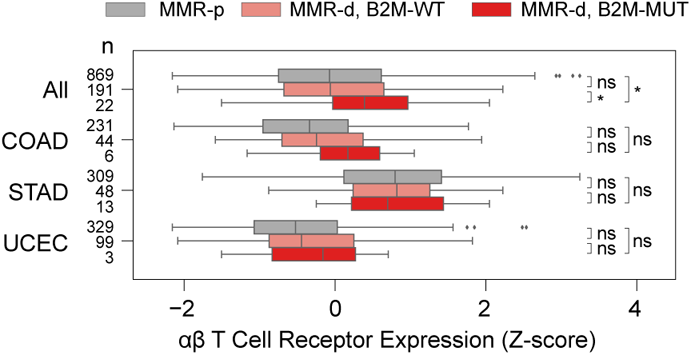
Association of B2M mutation status with expression of αβ T cell receptors. Boxplot showing the RNA expression of αβ T cell receptors in MMR-p (gray), MMR-d B2MWT (pink), and MMR-d B2MMUT (high impact; red) cancers. Results are obtained on the TCGA COAD, STAD and UCEC cohorts, and are shown for all cohorts combined (All), and for each cohort separately. Boxes, whiskers, and dots indicate quartiles, 1.5 interquartile ranges, and outliers, respectively. P-values were calculated by Wilcoxon rank sum test. *P<0.05.

**Extended Data Fig. 2.**
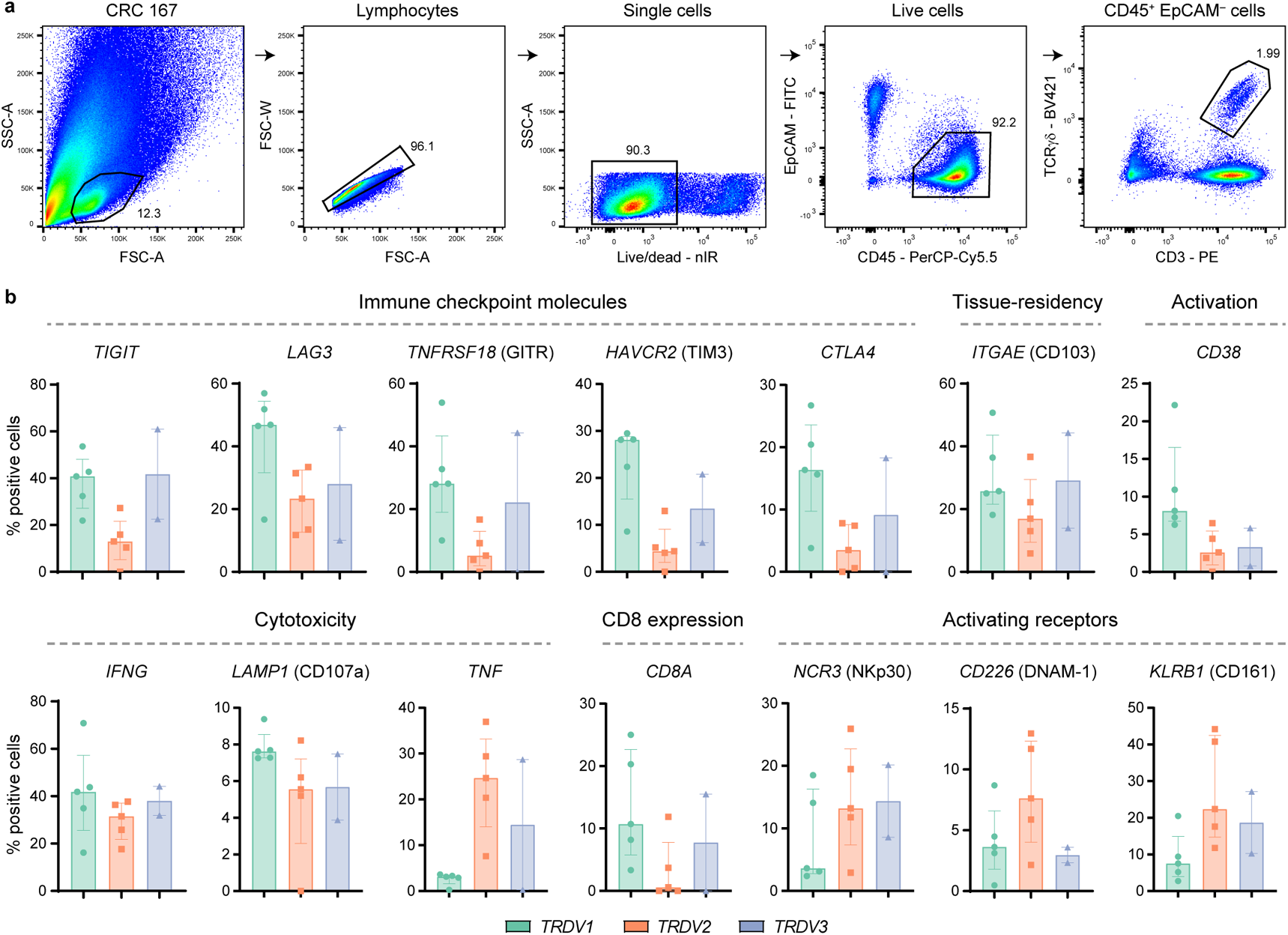
Characterization of γδ T cells from MMR-d colon cancers by single-cell RNA-sequencing. **a.** FACS gating strategy for single, live CD45+ EpCAM– CD3+ TCRγδ+ cells of a representative MMR-d colon cancer sample showing sequential gates with percentages. **b.** Frequencies of positive cells for selected genes across Vδ1 (n=1927), Vδ2 (n=860), and Vδ3 (n=506) cells as percentage of total γδ T cells from each MMR-d colon tumor (n=5) analyzed by single-cell RNA-sequencing. Vδ3 cells were present in two out of five colon cancers. Bars indicate median ± IQR. Each dot represents an individual sample.

**Extended Data Fig. 3.**
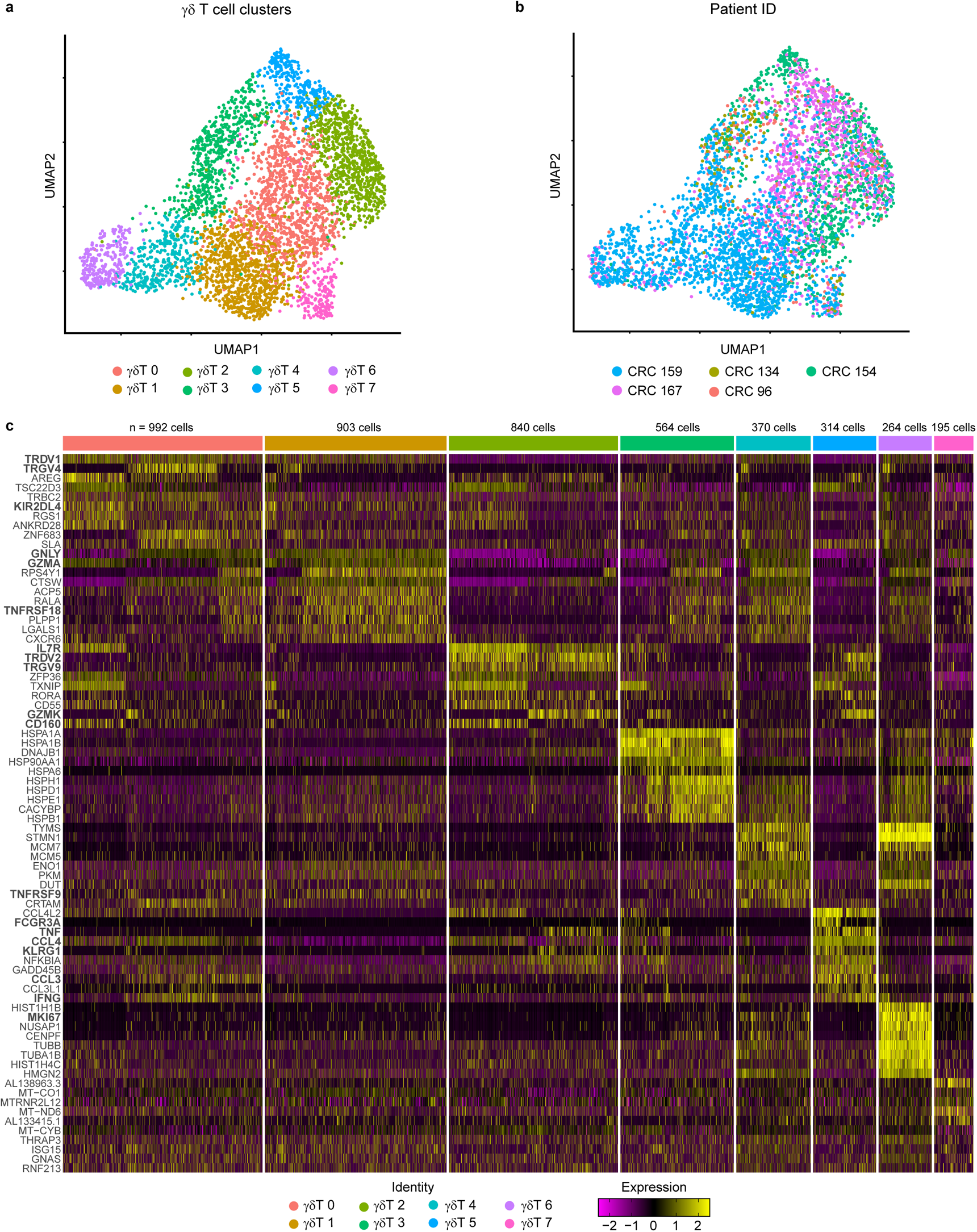
Distinct clusters of γδ T cells from MMR-d colon cancers by single-cell RNA-sequencing. **a.** UMAP embedding showing γδ T cells (n=4442) isolated from MMR-d colon cancers (n=5) analyzed by single-cell RNA-sequencing. Colors represent the functionally different γδ T cell clusters identified by graph-based clustering and non-linear dimensional reduction. Each dot represents a single cell. **b.** UMAP embedding of (a) colored by patient ID. Each dot represents a single cell. **c.** Heatmap showing the normalized single-cell gene expression value (z-score, purple-to-yellow scale) for the top 10 differentially expressed genes in each identified γδ T cell cluster.

**Extended Data Fig. 4.**
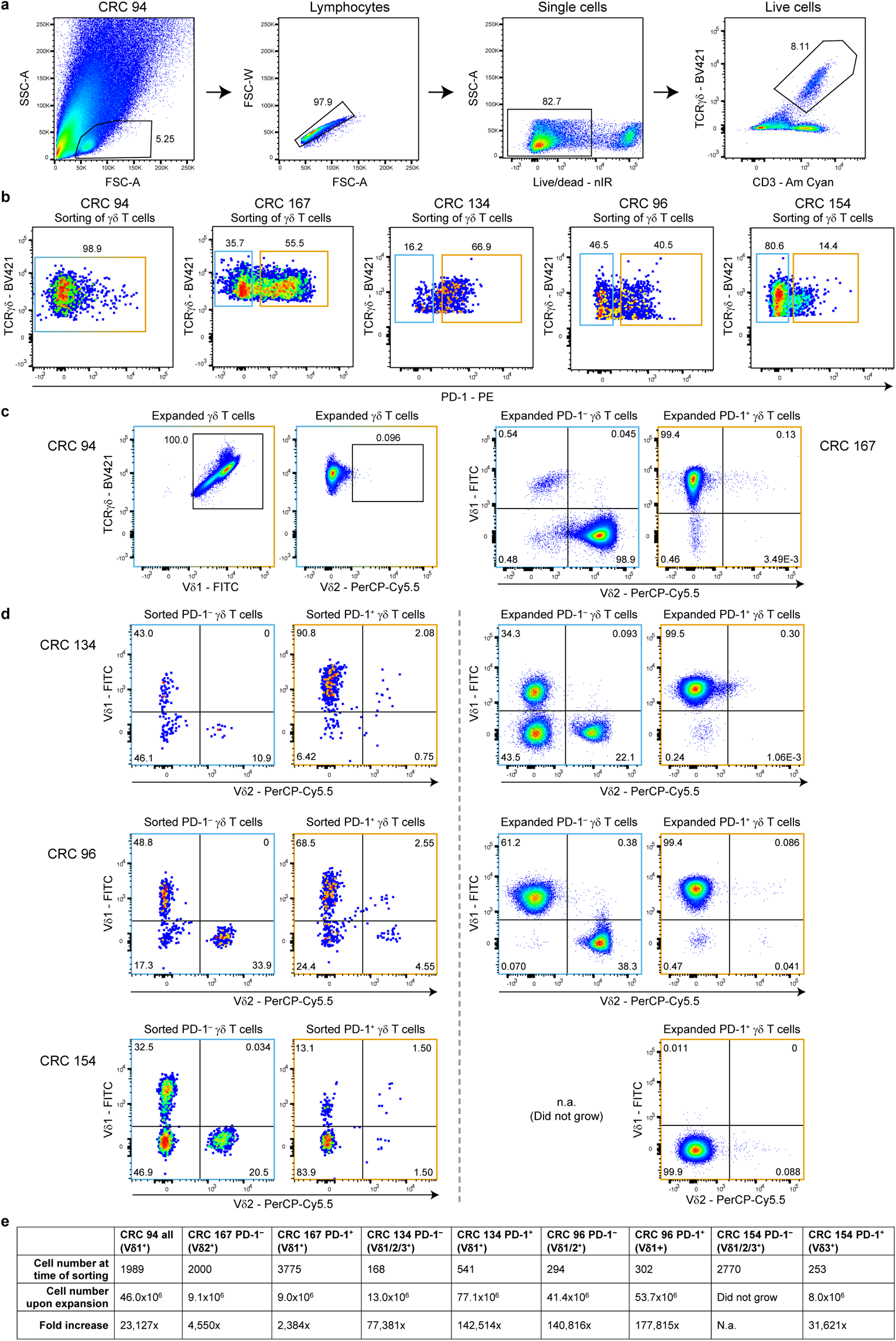
Sorting of PD-1+ and PD-1− γδ T cells from MMR-d colon cancers by FACS and their TCR Vδ chain usage. **a.** FACS gating strategy for single, live CD3+ TCRγδ+ cells of a representative MMR-d colon cancer sample showing sequential gates with percentages. **b.** Sorting of all γδ T cells from CRC94 (due to the low number of PD-1+ cells), and of PD-1– (blue squares) and PD-1+ (orange squares) γδ T cells from CRC167, CRC134, CRC96, and CRC154. Each dot is a single cell. **c.** TCR Vδ chain usage after expansion of γδ T cells from CRC94 and CRC167. Each dot is a single cell. **d.** TCR Vδ chain usage at the time of sorting (left panel) as well as after expansion of γδ T cells from CRC134, CRC96 and CRC154 (right panel). From CRC154, the PD-1– γδ T cells did not expand in culture. Each dot is a single cell. **e.** Table showing the number of γδ T cells isolated from colon cancers at the time of sorting versus 3-4 weeks after expansion, and the fold increase thereof.

**Extended Data Fig. 5.**
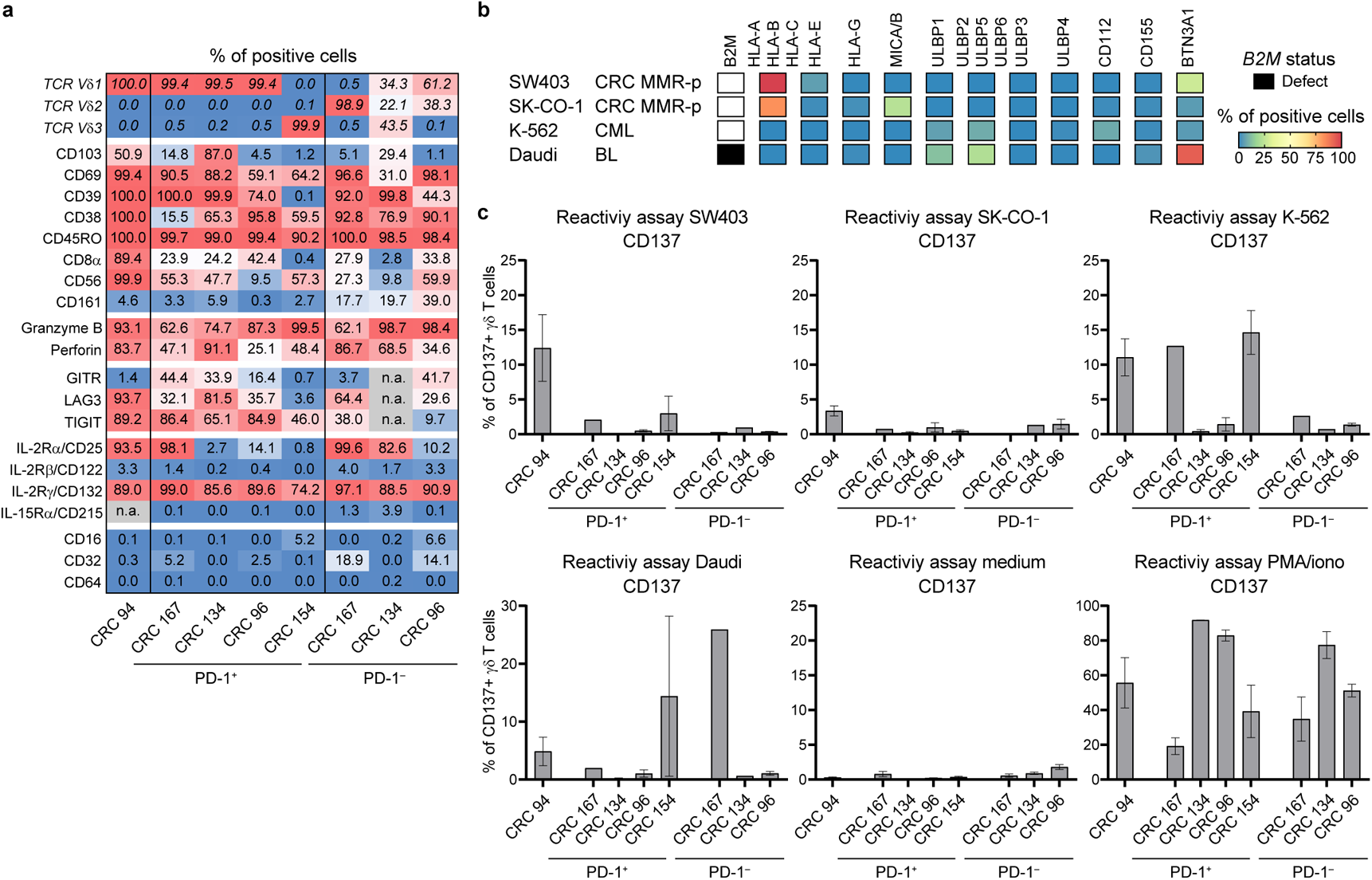
Reactivity of γδ T cells from MMR-d colon cancers towards cancer cell lines. **a.** Table showing the percentage of positive cells for different TCR Vδ chains (as in Fig. 3a), tissue-residency/activation markers, cytotoxic molecules, immune checkpoint molecules, cytokine receptors, and Fc receptors on expanded PD-1+ and PD-1– γδ T cells from MMR-d colon cancers (n=5) as percentage of total γδ T cells. **b.** Heatmap showing the B2M mutational status and surface expression of HLA class I, NKG2D ligands, DNAM-1 ligands, and butyrophilin on SW403, SK-CO-1, K-562, and Daudi cells. **c.** Bar plots showing the percentage of CD137-positive γδ T cells after 18h co-culture of PD-1+ and PD-1– γδ T cells from MMR-d colon cancers (n=5) with SW403, SK-CO-1, K-562, and Daudi cells. Medium was used as negative control and PMA/ionomycin as positive control. Bars indicate mean ± SEM. Data from two independent experiments (CRC94, CRC134, CRC154, CRC96), depending on availability of γδ T cells.

**Extended Data Fig. 6.**
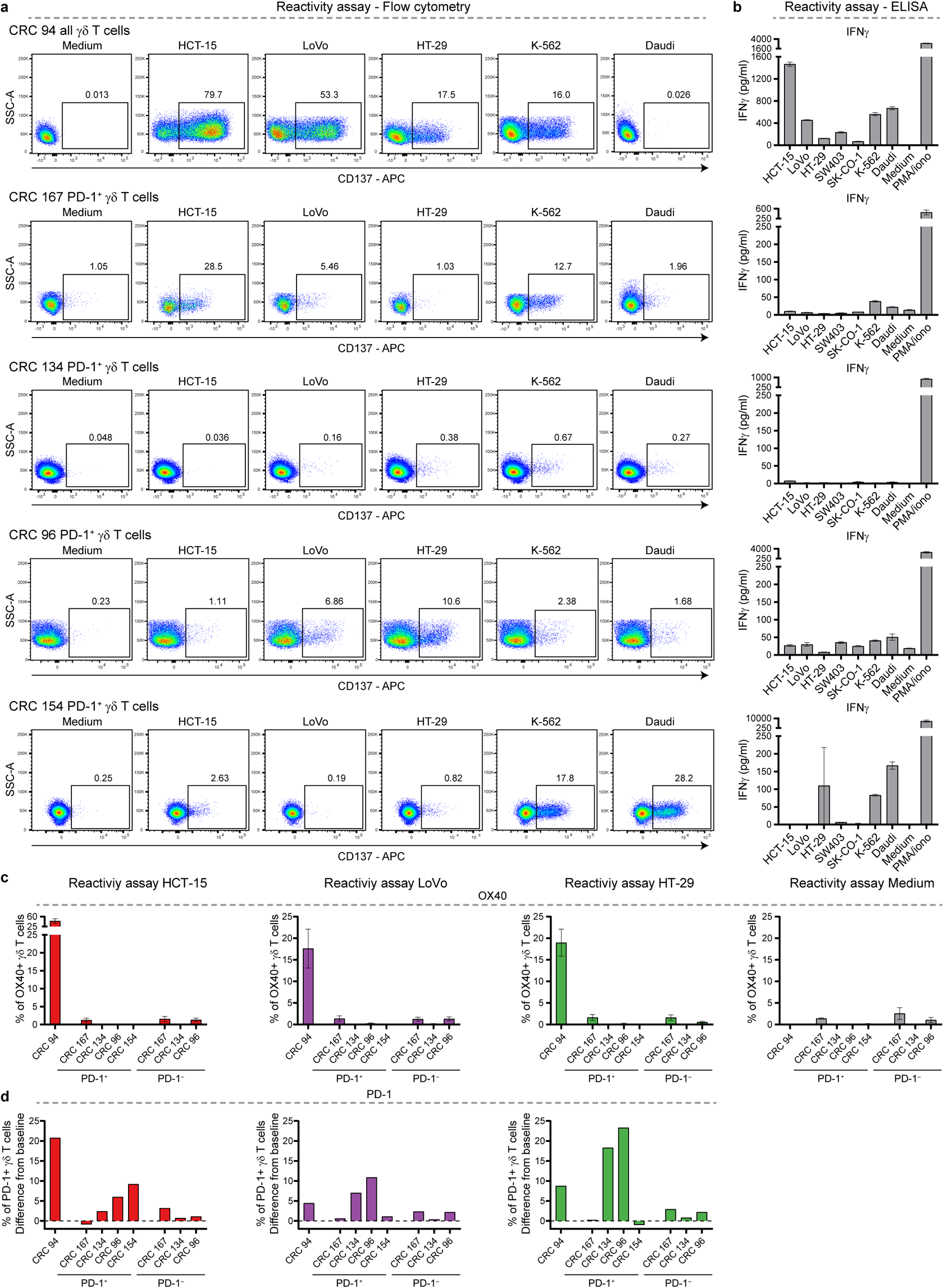
Surface expression of activation markers and secretion of IFNγ upon co-culture of γδ T cells from MMR-d colon cancers with cancer cell lines. **a.** Flow cytometry plots showing the expression of CD137 on PD-1+ γδ T cells after 18h co-culture with HCT-15, LoVo, HT-29, K-562, and Daudi cells as compared to medium only. Gates indicate percentage of positive γδ T cells. **b.** Bar plots showing the presence of IFNγ in the supernatant after 18h co-culture of PD-1+ γδ T cells with the cancer cell lines. Medium as negative control and PMA/ionomycin as positive control are included. Bars indicate mean ± SEM of triplicates. **c.** Bar plots showing the percentage of OX40-positive γδ T cells after 18h co-culture of PD-1+ and PD-1– γδ T cells from MMR-d colon cancers (n=5) with HCT-15, LoVo, and HT-29 cells. Bars indicate mean ± SEM. Data from four (CRC94), three (CRC167, CRC96), or two (CRC134, CRC154) independent experiments, depending on availability of γδ T cells. **d.** Bar plots showing the expression of PD-1 on γδ T cells as difference from baseline (medium) condition after 18h co-culture of PD-1+ and PD-1– γδ T cells from MMR-d colon cancers (n=5) with HCT-15, LoVo, and HT-29 cells.

**Extended Data Fig. 7.**
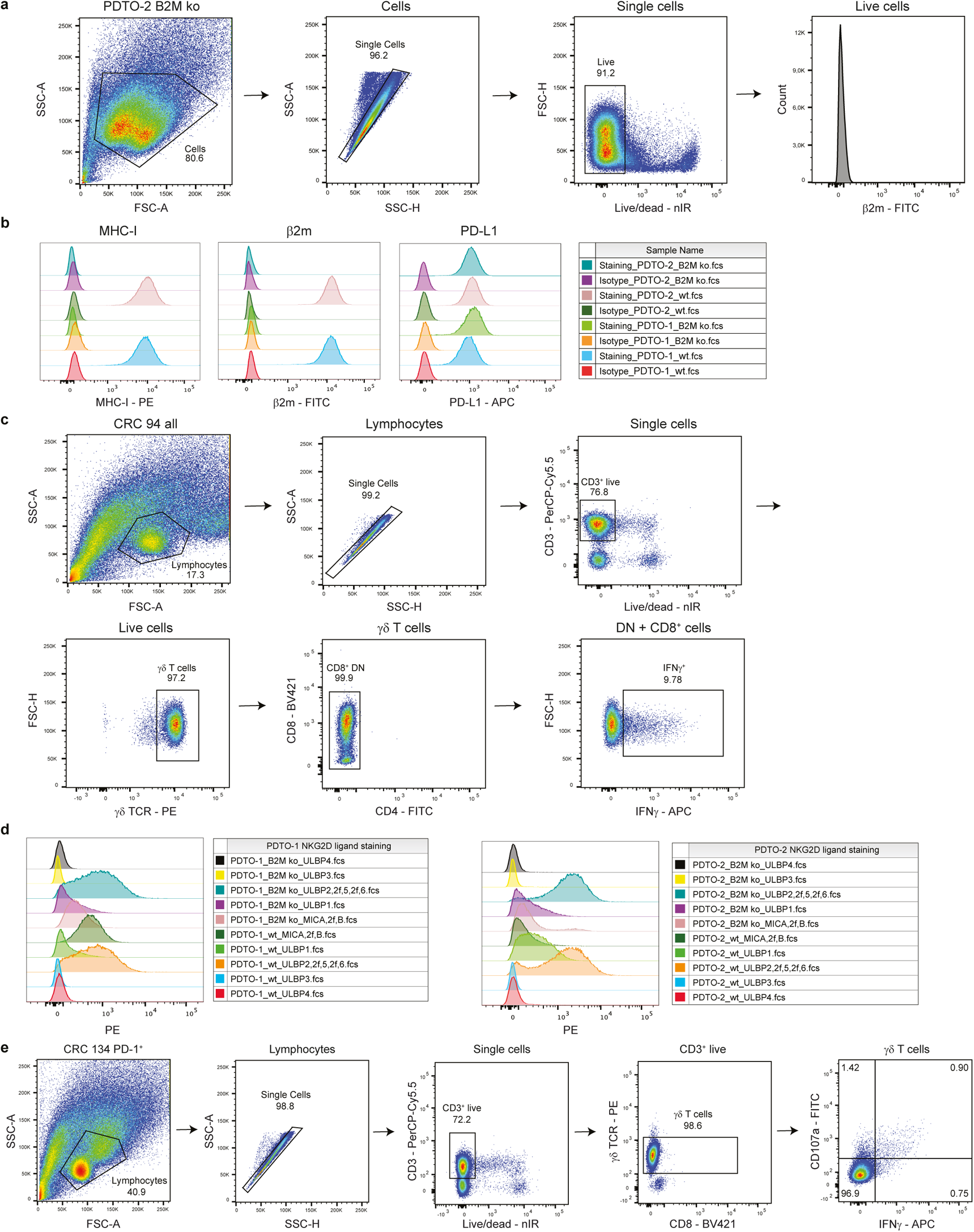
Tumor organoid characterization and reactivity assay readout. **a.** Flow cytometry gating strategy on PDTO cells for analysis of surface staining. Selected cells were gated on single, live cells before quantification of staining signal. **b.** Histogram representation and count for surface staining of MHC-I, PD-L1, and β2m expression on two PDTO lines B2MWT and B2MKO after IFNγ pre-stimulation. Staining with isotype antibodies for each fluorochrome (PE, APC and FITC) were included as negative control. **c.** Flow cytometry gating strategy on γδ T cell samples for analysis of intracellular staining to test antitumor reactivity upon PDTO stimulation. Lymphocyte population was further gated on single cells, live and CD3+ cells, γδ TCR+ cells and CD8+ as well as CD8–CD4– cells. Reactivity of the sample was based on IFNγ+ cells of the selected population. **d.** Histogram representation and count for surface staining of NKG2D ligands MICA/B, ULBP1, ULBP2/5/6, ULBP3, and ULBP4 on two PDTO lines B2MWT and B2MKO after IFNγ pre-stimulation. **e.** Flow cytometry gating strategy on γδ T cell samples for analysis of intracellular staining after stimulation with PDTOs in the presence of NKG2D ligand blocking. Lymphocyte population was further gated on single cells, live and CD3+ cells, followed by γδ TCR+ and CD8+ as well as CD8– cells. Reactivity of final population was based on IFNγ+ or CD107a+ cells.

**Extended Data Fig. 8.**
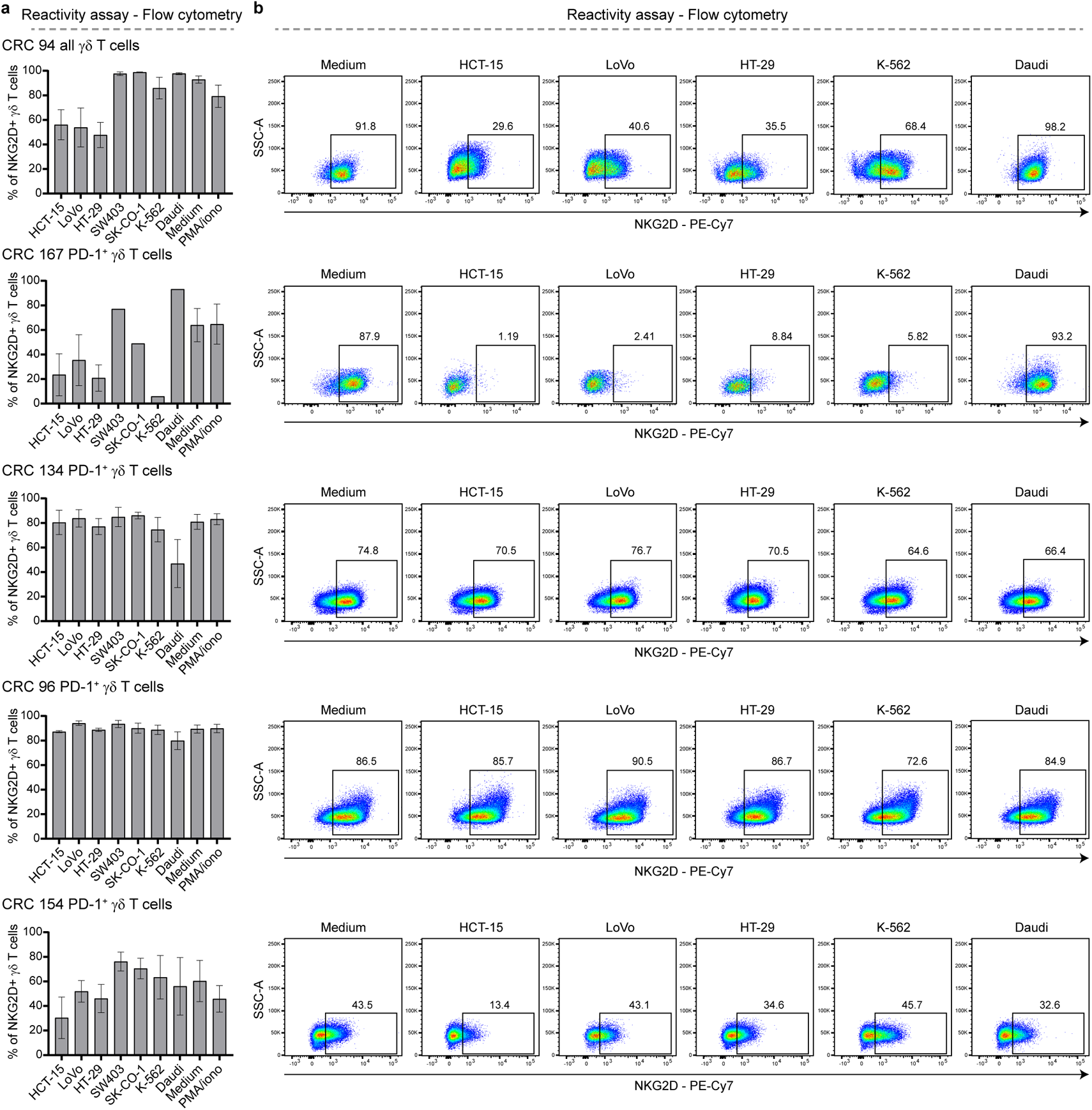
Surface expression of activating receptor NKG2D by PD-1+ γδ T cells from MMR-d colon cancers upon co-culture with cancer cell lines. **a.** Bar plots showing the expression of NKG2D on PD-1+ γδ T cells from MMR-d colon cancers (n=5) after 18h co-culture of PD-1+ γδ T cells with the cancer cell lines. Medium as negative control and PMA/ionomycin as positive control are included. Bars indicate mean ± SEM. Data from four (CRC94), three (CRC167, CRC96), or two (CRC134, CRC154) independent experiments, depending on availability of γδ T cells. **b.** Flow cytometry plots showing the expression of NKG2D on PD-1+ γδ T cells after 18h co-culture with HCT-15, LoVo, HT-29, K-562, and Daudi cells as compared to medium only. Gates indicate percentage of positive γδ T cells.

**Extended Data Fig. 9.**
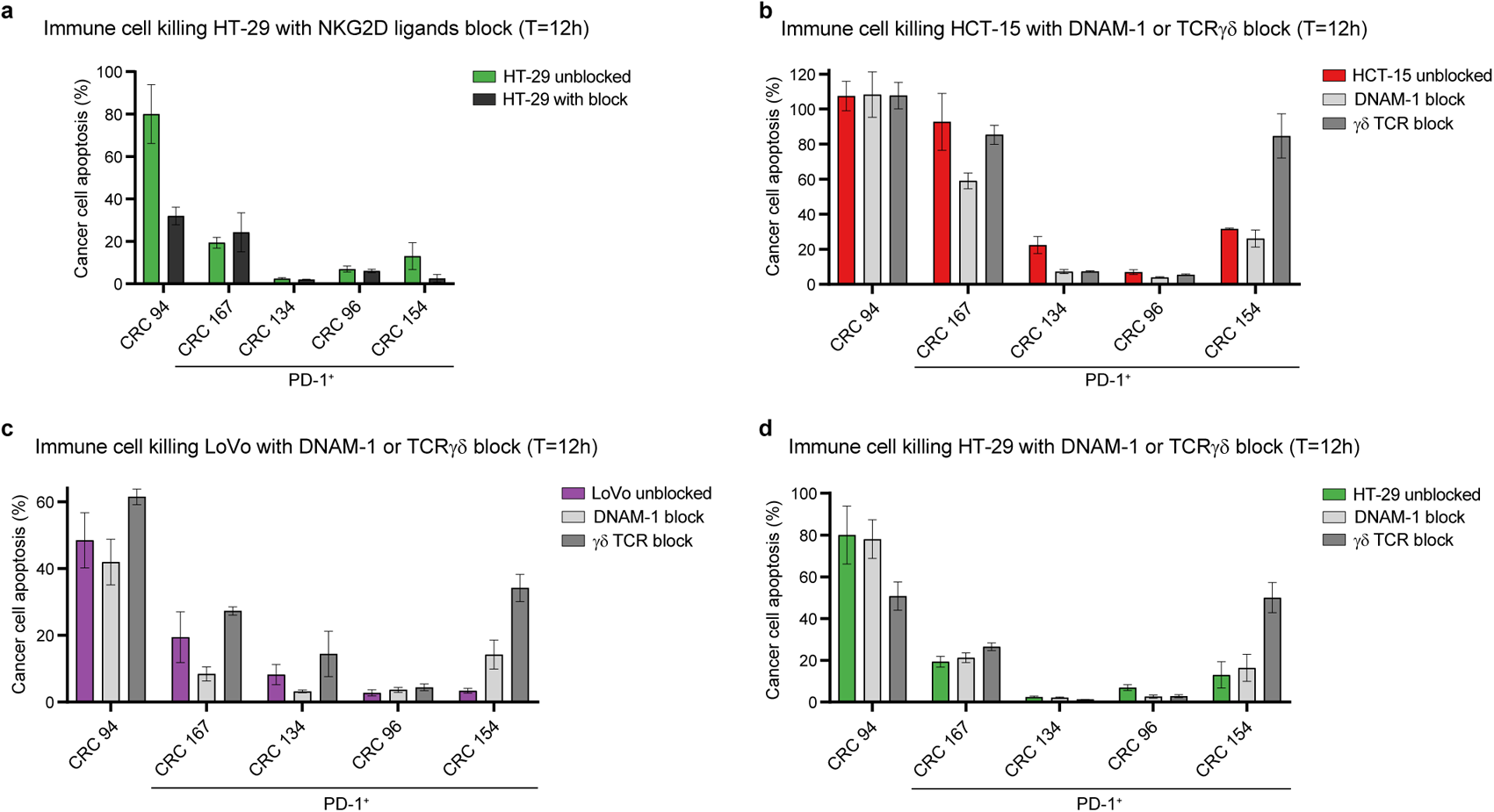
Killing of cancer cell lines by PD-1+ γδ T cells from MMR-d colon cancers in the presence of NKG2D ligand, DNAM-1, or γδ TCR blocking. **a.** Bar plots showing the quantification of killing of HT-29 cells by γδ T cells from MMR-d colon cancers (n=5) in the presence of blocking antibodies for NKG2D ligands as compared to the unblocked condition after 12h co-culture. Bars indicate mean ± SEM of two wells with two images/well. **b.** Bar plots showing the quantification of killing of HCT-15 cells by γδ T cells from MMR-d colon cancers (n=5) in the presence of DNAM-1 or γδ TCR blocking antibodies as compared to the unblocked condition after 12h co-culture. Bars indicate mean ± SEM of two wells with two images/well. **c.** As (b), but for LoVo cells. **d.** As (b), but for HT-29 cells.

**Extended Data Fig. 10.**
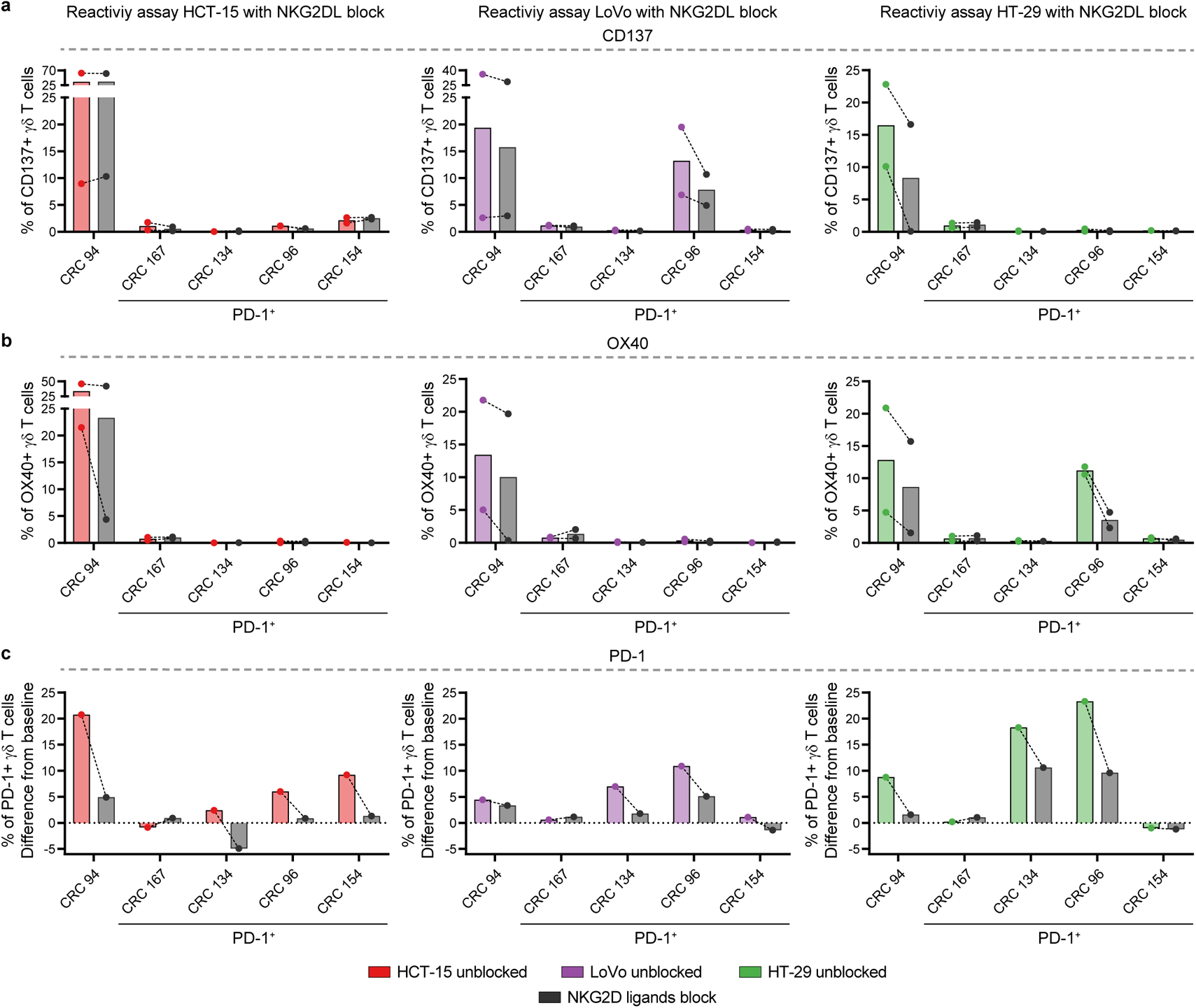
Reactivity towards cancer cell lines by PD-1+ γδ T cells from MMR-d colon cancers in the presence of NKG2D ligand blocking antibodies. **a.** Bar plots showing the percentage of CD137-positive γδ T cells after 18h co-culture of PD-1+ γδ T cells from MMR-d colon cancers (n=5) with HCT-15, LoVo, and HT-29 cells in the presence of blocking antibodies for NKG2D ligands. Lines indicate similar experiments. Data from two independent experiments. **b.** Bar plots showing the percentage of OX40-positive γδ T cells after 18h co-culture of PD-1+ γδ T cells from MMR-d colon cancers (n=5) with HCT-15, LoVo, and HT-29 cells in the presence of blocking antibodies for NKG2D ligands. Lines indicate similar experiments. Data from two independent experiments. **c.** Bar plots showing the expression of PD-1 on γδ T cells as difference from baseline (medium) condition after 18h co-culture of PD-1+ γδ T cells from MMR-d colon cancers (n=5) with HCT-15, LoVo, and HT-29 cells in the presence of blocking antibodies for NKG2D ligands. Lines indicate similar experiments.

**Extended Data Fig. 11.**
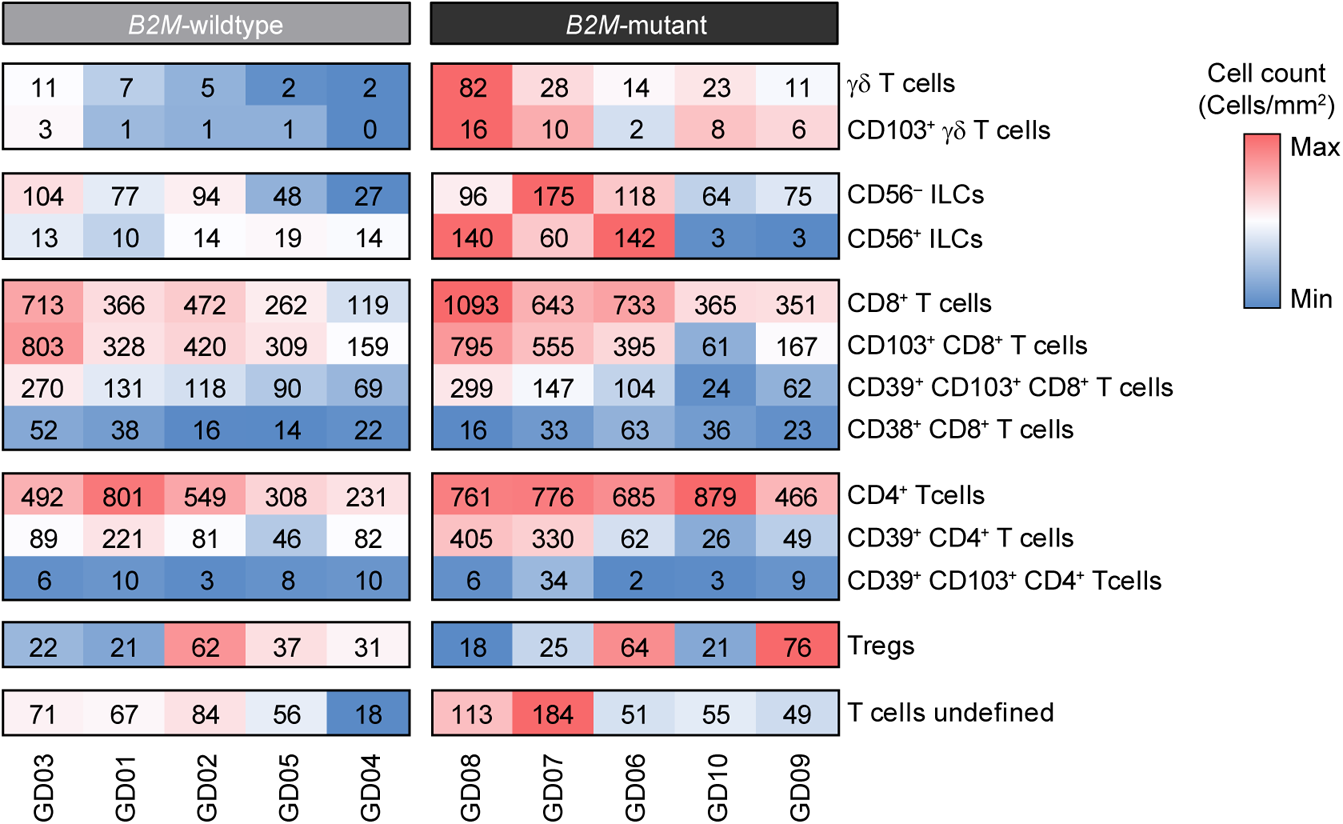
Distribution of immune cell populations in B2M-wildtype and B2M-mutant colon cancers upon immune checkpoint blockade (ICB) by imaging mass cytometry. Heatmap showing cell counts (cells/mm2) of different immune cell phenotypes from the imaging mass cytometric detection of B2MWT (HLA class I-positive, n=5) and B2MMUT (HLA class I-negative, n=5) MMR-d colon cancers upon ICB treatment. Hierarchical clustering was performed on the samples within the two groups. Color bar is scaled per major immune lineage.

**Extended Data Table 1.** Characteristics of clinical samples from 17 patients with MMR-deficient colon cancer.

| Patient ID | Tumor location | MMR status | HLA class I status | β2m status | Subgroup |
| --- | --- | --- | --- | --- | --- |
| CRC 1 | Flexura lienalis | MMR-deficient | Positive | Positive | HLA+ |
| CRC 2 | Coecum | MMR-deficient | Defect | Mut pattern | HLA defect |
| CRC 3 | Flexura hepatica | MMR-deficient | Positive | Positive | HLA+ |
| CRC 6 | Descendens | MMR-deficient | Defect | Defect | B2M defect |
| CRC 19 | Ascendens | MMR-deficient | Defect | Defect | B2M defect |
| CRC 52 | Ascendens | MMR-deficient | Defect | Mut pattern | HLA defect |
| CRC 67 | Transversum | MMR-deficient | Positive | Positive | HLA+ |
| CRC 84 | Sigmoid | MMR-deficient | Defect | Defect | B2M defect |
| CRC 94 # | Sigmoid | MMR-deficient | Defect | Positive | HLA defect |
| CRC 96* # | Transversum | MMR-deficient | Defect | Defect | B2M defect |
| CRC 102 | Ascendens | MMR-deficient | Defect | Positive | HLA defect |
| CRC 134* # | Ascendens | MMR-deficient | Defect | Defect | B2M defect |
| CRC 154* # | Flexura hepatica | MMR-deficient | Positive | Positive | HLA+ |
| CRC 159* | Coecum | MMR-deficient | Defect | Mut pattern | HLA defect |
| CRC 167* # | Ascendens | MMR-deficient | Defect | Mut pattern | HLA defect |
| CRC 177 | Flexura hepatica | MMR-deficient | Defect | Positive | HLA defect |
| CRC 195 | Flexura hepatica | MMR-deficient | Defect | Positive | HLA defect |

\* Used for single-cell RNA-sequencing experiments, # Used for cell culturing experiments. B2M, β2-microglobulin; MMR, mismatch repair.

**Extended Data Table 2.**
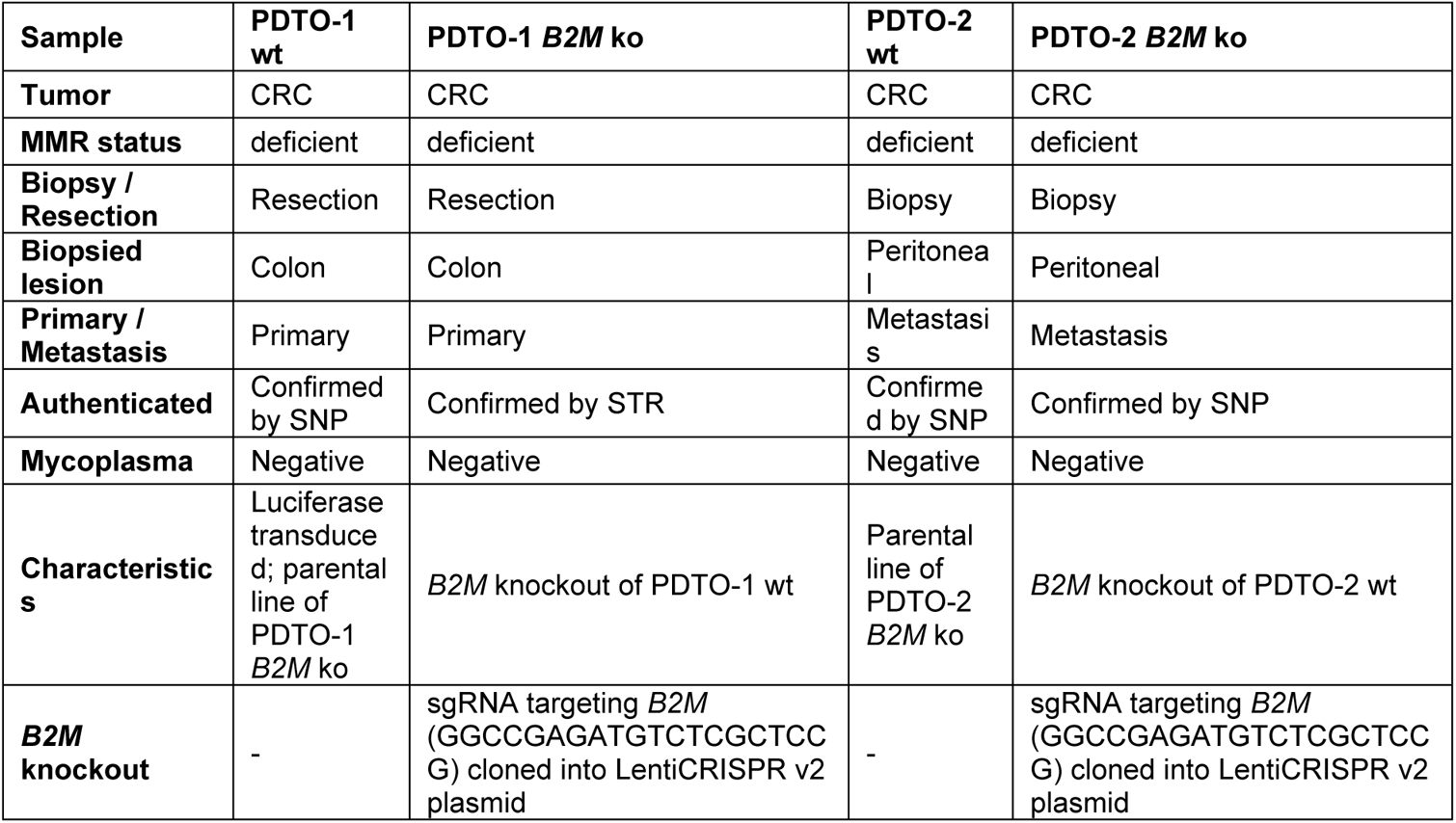
Characteristics of patient-derived organoids from MMR-deficient colorectal cancer.

**Extended Data Table 3.** Antibodies used for imaging mass cytometry of colon cancers.

| Antibody | Metal | Clone | Supplier | Catalog number | Lot number | Incubation time (temperature) | Dilution |
| --- | --- | --- | --- | --- | --- | --- | --- |
| β-catenin | 89Y | D10A8 | CST | 8480BF | 8 | Overnight (4°C) | 1:100 |
| CD103 | 168 Er | EPR4166(2) | Abcam | ab221210 | GR3355784-7 | 5h (RT) | 1:50 |
| CD11b | 144 Nd | D6X1N | CST | 49420BF | 4 | 5h (RT) | 1:100 |
| CD11c | 176 Yb | EP1347Y | Abcam | ab216655 | GR3357092-9 | 5h (RT) | 1:100 |
| CD14 | 163 Dy | D7A2T | CST | 56082BF | 2 | 5h (RT) | 1:100 |
| CD15 | 171 Yb | MC480 | CST | 4744BF | 5 | Overnight (4°C) | 1:100 |
| CD163 | 173 Yb | EPR14643-36 | Abcam | 93498BF | Not available | 5h (RT) | 1:50 |
| CD20 | 142 Nd | E7B7T | CST | 48750BF | 9179056 | Overnight (4°C) | 1:100 |
| CD204 | 164 Dy | J5HTR3 | Thermo Fisher Scientific | 14-9054-95 | 4338161 | 5h (RT) | 1:50 |
| CD3 | 153 Eu | EP449E | Abcam | ab271850 | GR3341846-3 | Overnight (4°C) | 1:50 |
| CD31 | 147 Sm | 89C2 | CST | 3528BF | Not available | Overnight (4°C) | 1:100 |
| CD38 | 169 Tm | EPR4106 | Abcam | ab226034 | GR3378690-1 | Overnight (4°C) | 1:100 |
| CD39 | 157 Gd | EPR20627 | Abcam | ab236038 | GR3274485-6 | 5h (RT) | 1:100 |
| CD4* | 145 Nd | EPR6855 | Abcam | ab181724 | GR3285644-10 | Overnight (RT) | 1:100 |
| CD45 | 149 Sm | D9M8I | CST | 13917BF | 11 | Overnight (4°C) | 1:50 |
| CD45RO | 165 Ho | UCHL1 | CST | 55618BF | 2 | Overnight (4°C) | 1:100 |
| CD56 | 167 Er | E7X9M | CST | 99746BF | 2 | 5h (RT) | 1:100 |
| CD57 | 151 Eu | HNK-1 / Leu-7 | Abcam | ab269781 | GR3373313-3 | Overnight (4°C) | 1:100 |
| CD68 | 143 Nd | D4B9C | CST | 76437BF | 2 | Overnight (4°C) | 1:100 |
| CD7 | 174 Yb | EPR4242 | Abcam | ab230834 | Not available | 5h (RT) | 1:100 |
| CD8α | 146 Nd | D8A8Y | CST | 85336BF | Not available | 5h (RT) | 1:50 |
| Cleaved caspase-3 | 172 Yb | 5A1E | CST | 9664BF | 24 | 5h (RT) | 1:100 |
| D2-40 | 166 Er | D2-40 | BioLegend | 916606 | B316467 | Overnight (4°C) | 1:100 |
| FOXP3 | 159 Tb | D608R | CST | 12653BF | 8 | Overnight (4°C) | 1:50 |
| Granzyme B | 150 Nd | D6E9W | CST | 46890BF | 3 | 5h (RT) | 1:100 |
| Histone H3 | 209 | D1H2 | CST | 4499BF | Not available | Overnight (4°C) | 1:50 |
| HLA-DR | 141 Pr | TAL 1B5 | Abcam | ab176408 | GR3384096-1 | 5h (RT) | 1:100 |
| ICOS | 161 Dy | D1K2T™ | CST | 89601BF | 4 | 5h (RT) | 1:50 |
| IDO | 162 Dy | D5J4E™ | CST | 86630BF | 7 | Overnight (4°C) | 1:100 |
| Ki-67 | 152 Sm | 8D5 | CST | 9449BF | 11 | Overnight (4°C) | 1:100 |
| LAG-3 | 155<br>Gd | D2G40™ | CST | 15372BF | Not available | 5h (RT) | 1:50 |
| p16ink4a | 175<br>Lu | D3W8G | CST | 92803BF | 2 | Overnight (4°C) | 1:100 |
| Pan-keratin | 198<br>Pt | C11 and<br>AE1/AE3 | CST /<br>BioLegend | 4545BF /<br>914204 | 12 /<br>B302316 | Overnight (4°C) | 1:50 |
| PD-1 | 160<br>Gd | D4W2J | CST | 86163BF | 7 | 5h (RT) | 1:50 |
| PD-L1 | 156<br>Gd | E1L3N<sup>R</sup> | CST | 13684BF | 17 | Overnight (4°C) | 1:50 |
| T-bet | 170<br>Er | 4B10 | BioLegend | 644825 | B298378 | 5h (RT) | 1:50 |
| TCRδ* | 148<br>Nd | H41 | Santa Cruz | sc-100289 | D3021 | Overnight (RT) | 1:50 |
| TGFβ | 115<br>In | TB21 | Thermo<br>Fisher<br>Scientific | MA5-16949 | 151471 | 5h (RT) | 1:100 |
| TIM-3 | 154<br>Sm | D5D5R™ | CST | 45208BF | 9 | 5h (RT) | 1:100 |
| Vimentin | 194<br>Pt | D21H3 | CST | 5741BF | 9 | Overnight (4°C) | 1:50 |
| VISTA | 158<br>Gd | D1L2G™ | CST | 64953BF | 7 | 5h (RT) | 1:100 |

\*Detection with metal-conjugated secondary antibodies. CST, Cell Signaling Technology.

**Extended Data Table 4.** Antibodies used for immunophenotyping of γδ T cells by flow cytometry.

| Antibody | Fluorochrome | Clone | Supplier | Catalog number | Lot number | Dilution |
| --- | --- | --- | --- | --- | --- | --- |
| CD16 | PE | B73.1 | BD Biosciences | 332779 | 9045985 | 1:60 |
| CD103 | FITC | Ber-ACT8 | BD Biosciences | 550259 | 2332847 | 1:10 |
| CD122/IL-2Rβ | BV421 | TU27 | BioLegend | 339010 | B313155 | 1:20 |
| CD132/IL-2Rγ | APC | TUGh4 | BioLegend | 338608 | B293032 | 1:80 |
| CD161 | BV605 | DX12 | BD Biosciences | 563863 | 7030586 | 1:20 |
| CD25/IL-2Rα | PE-Cy7 | M-A251 | BD Biosciences | 557741 | 9301660 | 1:25 |
| CD215/IL-15Rα | PE | JM7A4 | BioLegend | 330208 | B265801 | 1:80 |
| CD226/DNAM-1 | BV510 | DX11 | BD Biosciences | 742494 | 9203072 | 1:150 |
| CD3 | Am Cyan | SK7 | BD Biosciences | 339186 | 9161745 | 1:20 |
| CD32 | APC | FLI8.26 | BD Biosciences | 559769 | 184743 | 1:20 |
| CD38 | PE-Cy7 | HIT2 | eBioscience | 25-0389-42 | 4319912 | 1:200 |
| CD39 | APC | A1 | BioLegend | 328210 | B249211 | 1:60 |
| CD45RA | FITC | L48 | BD Biosciences | 335039 | 8227525 | 1:30 |
| CD45RA | PE-Dazzle594 | HI100 | Sony | 2120730 | 126470 | 1:20 |
| CD45RO | PerCP-Cy5.5 | UCHL1 | Sony | 2121110 | 138351 | 1:20 |
| CD56 | APC-R700 | NCAM16.2 | BD Biosciences | 565139 | 5251693 | 1:150 |
| CD64 | FITC | 10.1 | BD Biosciences | 555527 | 58058 | 1:20 |
| CD69 | PerCP-Cy5.5 | FN50 | BioLegend | 310925 | B266970 | 1:200 |
| CD8α | BV605 | SK1 | BD Biosciences | 564115 | 7092 | 1:100 |
| CD94 | BV605 | HP-3D9 | BD Biosciences | 743950 | 7138571 | 1:200 |
| GITR | PE | 108-17 | BioLegend | 371204 | B244963 | 1:50 |
| Granzyme B* | PE | GB11 | eBiosciences | 12-8899-41 | 1928380 | 1:50 |
| KIR2DL1 | PE | HP3-E4 | BD Biosciences | 556063 | 86798 | 1:20 |
| KIR2DL1/S1 | PE | EB6 | Beckman Coulter | A09778 | 12 | 1:50 |
| KIR2DL2/L3/S2 | PE | GL183 | Beckman Coulter | IM2278U | 200051 | 1:50 |
| KIR2DL4 | PE | 181703 | R&D Systems | FAB2238P | AAHO0209081 | 1:10 |
| KIR2DS4 | PE | FES172 | Beckman Coulter | IM3337 | 200037 | 1:40 |
| KIR3DL1 | PE | DX9 | BD Biosciences | 555967 | 121769 | 1:80 |
| KIR3DL1/S1 | PE | Z27 | Beckman Coulter | IM3292 | 200044 | 1:20 |
| KIR3DL2 | PE | #539304 | R&D Systems | FAB2878P | ADBO0217051 | 1:10 |
| LAG3 | PE-Cy7 | 11C3C65 | BioLegend | 369309 | B289009 | 1:100 |
| NKG2A | APC | z199 | Beckman Coulter | A60797 | 200046 | 1:30 |
| NKG2C | PE | 134591 | Beckman Coulter | FAB138P | LCN0818011 | 1:20 |
| NKG2D | PE-Cy7 | 1D11 | BD Biosciences | 562365 | 9045733 | 1:300 |
| NKp44 | APC | P44-8 | BioLegend | 325109 | B160899 | 1:20 |
| NKp46 | PE | 9E2 | BioLegend | 331907 | B150121 | 1:20 |
| PD-1 | PE | MIH4 | eBioscience | 12-9969-42 | 1952441 | 1:30 |
| Perforin* | PE-Cy7 | dG9 | BioLegend | 308125 | B215704 | 1:20 |
| TCR$\gamma\delta$ | BV421 | 11F2 | BD Biosciences | 744870 | 9340519 | 1:80 |
| TCR$\gamma\delta$ | BV650 | 11F2 | BD Biosciences | 745359 | 7222894 | 1:40 |
| TCR Vδ1 | FITC | TS8.2 | Invitrogen | TCR2730 | UH286015 | 1:50 |
| TCR Vδ2 | PerCP-Cy5.5 | B6 | BioLegend | 331424 | B279957 | 1:200 |
| TIGIT | APC | 1D11 | BD Biosciences | 562365 | 9045733 | 1:300 |
| Live/dead | nIR | n.a. | Life Technologies | L10119 | 1808830 | 1:1000 |

\*Detected intracellularly

## Notes

### Competing Interest Statement

The authors have declared no competing interest.

## References

1. Ionov, Y., Peinado, M. A., Malkhosyan, S., Shibata, D. & Perucho, M. Ubiquitous somatic mutations in simple repeated sequences reveal a new mechanism for colonic carcinogenesis. Nature 363, 558–561, doi:10.1038/363558a0 (1993).

2. Germano, G. et al. Inactivation of DNA repair triggers neoantigen generation and impairs tumour growth. Nature 552, 116–120, doi:10.1038/nature24673 (2017).

3. Bicknell, D. C., Kaklamanis, L., Hampson, R., Bodmer, W. F. & Karran, P. Selection for beta 2-microglobulin mutation in mismatch repair-defective colorectal carcinomas. Current biology : CB 6, 1695–1697, doi:10.1016/s0960-9822(02)70795-1 (1996).

4. Dierssen, J. W. et al. HNPCC versus sporadic microsatellite-unstable colon cancers follow different routes toward loss of HLA class I expression. BMC Cancer 7, 33, doi:10.1186/1471-2407-7-33 (2007).

5. Kloor, M. et al. Immunoselective pressure and human leukocyte antigen class I antigen machinery defects in microsatellite unstable colorectal cancers. Cancer Res 65, 6418–6424, doi:10.1158/0008-5472.can-05-0044 (2005).

6. Ijsselsteijn, M. E. et al. Revisiting immune escape in colorectal cancer in the era of immunotherapy. Br J Cancer 120, 815–818, doi:10.1038/s41416-019-0421-x (2019).

7. Middha, S. et al. Majority of B2M-Mutant and -Deficient Colorectal Carcinomas Achieve Clinical Benefit From Immune Checkpoint Inhibitor Therapy and Are Microsatellite Instability-High. JCO precision oncology 3, doi:10.1200/po.18.00321 (2019).

8. Le, D. T. et al. Mismatch repair deficiency predicts response of solid tumors to PD-1 blockade. Science 357, 409–413, doi:10.1126/science.aan6733 (2017).

9. Overman, M. J. et al. Nivolumab in patients with metastatic DNA mismatch repair-deficient or microsatellite instability-high colorectal cancer (CheckMate 142): an open-label, multicentre, phase 2 study. The Lancet. Oncology 18, 1182–1191, doi:10.1016/s1470-2045(17)30422-9 (2017).

10. Overman, M. J. et al. Durable Clinical Benefit With Nivolumab Plus Ipilimumab in DNA Mismatch Repair-Deficient/Microsatellite Instability-High Metastatic Colorectal Cancer. Journal of clinical oncology : official journal of the American Society of Clinical Oncology 36, 773–779, doi:10.1200/jco.2017.76.9901 (2018).

11. Chalabi, M. et al. Neoadjuvant immunotherapy leads to pathological responses in MMR-proficient and MMR-deficient early-stage colon cancers. Nat Med 26, 566–576, doi:10.1038/s41591-020-0805-8 (2020).

12. Dolcetti, R. et al. High prevalence of activated intraepithelial cytotoxic T lymphocytes and increased neoplastic cell apoptosis in colorectal carcinomas with microsatellite instability. Am J Pathol 154, 1805–1813, doi:10.1016/s0002-9440(10)65436-3 (1999).

13. Tumeh, P. C. et al. PD-1 blockade induces responses by inhibiting adaptive immune resistance. Nature 515, 568–571, doi:10.1038/nature13954 (2014).

14. Taube, J. M. et al. Association of PD-1, PD-1 ligands, and other features of the tumor immune microenvironment with response to anti-PD-1 therapy. Clin Cancer Res 20, 5064–5074, doi:10.1158/1078-0432.Ccr-13-3271 (2014).

15. Groh, V. et al. Human lymphocytes bearing T cell receptor gamma/delta are phenotypically diverse and evenly distributed throughout the lymphoid system. J Exp Med 169, 1277–1294, doi:10.1084/jem.169.4.1277 (1989).

16. Silva-Santos, B., Serre, K. & Norell, H. γδ T cells in cancer. Nat Rev Immunol 15, 683–691, doi:10.1038/nri3904 (2015).

17. Halary, F. et al. Control of self-reactive cytotoxic T lymphocytes expressing gamma delta T cell receptors by natural killer inhibitory receptors. Eur J Immunol 27, 2812–2821, doi:10.1002/eji.1830271111 (1997).

18. de Vries, N. L. et al. High-dimensional cytometric analysis of colorectal cancer reveals novel mediators of antitumour immunity. Gut 69, 691–703, doi:10.1136/gutjnl-2019-318672 (2020).

19. Danaher, P. et al. Gene expression markers of Tumor Infiltrating Leukocytes. J Immunother Cancer 5, 18, doi:10.1186/s40425-017-0215-8 (2017).

20. Duhen, T. et al. Co-expression of CD39 and CD103 identifies tumor-reactive CD8 T cells in human solid tumors. Nat Commun 9, 2724, doi:10.1038/s41467-018-05072-0 (2018).

21. Kwon, M. et al. Determinants of Response and Intrinsic Resistance to PD-1 Blockade in Microsatellite Instability-High Gastric Cancer. Cancer Discov, doi:10.1158/2159-8290.Cd-21-0219 (2021).

22. Wu, P. et al. γδT17 cells promote the accumulation and expansion of myeloid­derived suppressor cells in human colorectal cancer. Immunity 40, 785–800, doi:10.1016/j.immuni.2014.03.013 (2014).

23. Lo Presti, E., Dieli, F. & Meraviglia, S. Tumor-Infiltrating γδ T Lymphocytes: Pathogenic Role, Clinical Significance, and Differential Programing in the Tumor Microenvironment. Front Immunol 5, 607, doi:10.3389/fimmu.2014.00607 (2014).

24. Maeurer, M. J. et al. Human intestinal Vdelta1+ lymphocytes recognize tumor cells of epithelial origin. J Exp Med 183, 1681–1696, doi:10.1084/jem.183.4.1681 (1996).

25. Mikulak, J. et al. NKp46-expressing human gut-resident intraepithelial Vδ1 T cell subpopulation exhibits high antitumor activity against colorectal cancer. JCI insight 4, doi:10.1172/jci.insight.125884 (2019).

26. Wu, D. et al. Ex vivo expanded human circulating Vδ1 γδT cells exhibit favorable therapeutic potential for colon cancer. Oncoimmunology 4, e992749, doi:10.4161/2162402x.2014.992749 (2015).

27. Siegers, G. M., Ribot, E. J., Keating, A. & Foster, P. J. Extensive expansion of primary human gamma delta T cells generates cytotoxic effector memory cells that can be labeled with Feraheme for cellular MRI. Cancer Immunol Immunother 62, 571–583, doi:10.1007/s00262-012-1353-y (2013).

28. Almeida, A. R. et al. Delta One T Cells for Immunotherapy of Chronic Lymphocytic Leukemia: Clinical-Grade Expansion/Differentiation and Preclinical Proof of Concept. Clin Cancer Res 22, 5795–5804, doi:10.1158/1078-0432.Ccr-16-0597 (2016).

29. van der Leun, A. M., Thommen, D. S. & Schumacher, T. N. CD8(+) T cell states in human cancer: insights from single-cell analysis. Nat Rev Cancer 20, 218–232, doi:10.1038/s41568-019-0235-4 (2020).

30. Groh, V. et al. Broad tumor-associated expression and recognition by tumor-derived gamma delta T cells of MICA and MICB. Proc Natl Acad Sci U S A 96, 6879–6884, doi:10.1073/pnas.96.12.6879 (1999).

31. Poggi, A. et al. Vdelta1 T lymphocytes from B-CLL patients recognize ULBP3 expressed on leukemic B cells and up-regulated by trans-retinoic acid. Cancer Res 64, 9172–9179, doi:10.1158/0008-5472.Can-04-2417 (2004).

32. Hause, R. J., Pritchard, C. C., Shendure, J. & Salipante, S. J. Classification and characterization of microsatellite instability across 18 cancer types. Nat Med 22, 1342–1350, doi:10.1038/nm.4191 (2016).

33. Cader, F. Z. et al. A peripheral immune signature of responsiveness to PD-1 blockade in patients with classical Hodgkin lymphoma. Nat Med 26, 1468–1479, doi:10.1038/s41591-020-1006-1 (2020).

34. Germano, G. et al. CD4 T cell dependent rejection of beta 2 microglobulin null mismatch repair deficient tumors. Cancer Discov, doi:10.1158/2159-8290.Cd-20-0987 (2021).

## Methods references

35. Thorsson, V. et al. The Immune Landscape of Cancer. Immunity 51, 411–412, doi:10.1016/j.immuni.2019.08.004 (2019).

36. Dobin, A. et al. STAR: ultrafast universal RNA-seq aligner. Bioinformatics (Oxford, England) 29, 15–21, doi:10.1093/bioinformatics/bts635 (2013).

37. Robinson, M. D., McCarthy, D. J. & Smyth, G. K. edgeR: a Bioconductor package for differential expression analysis of digital gene expression data. Bioinformatics (Oxford, England) 26, 139–140, doi:10.1093/bioinformatics/btp616 (2010).

38. Ritchie, M. E. et al. limma powers differential expression analyses for RNA-sequencing and microarray studies. Nucleic Acids Res 43, e47, doi:10.1093/nar/gkv007 (2015).

39. Law, C. W., Chen, Y., Shi, W. & Smyth, G. K. voom: Precision weights unlock linear model analysis tools for RNA-seq read counts. Genome Biol 15, R29, doi:10.1186/gb-2014-15-2-r29 (2014).

40. Davoli, T., Uno, H., Wooten, E. C. & Elledge, S. J. Tumor aneuploidy correlates with markers of immune evasion and with reduced response to immunotherapy. Science 355, doi:10.1126/science.aaf8399 (2017).

41. Virtanen, P. et al. SciPy 1.0: fundamental algorithms for scientific computing in Python. Nat Methods 17, 261–272, doi:10.1038/s41592-019-0686-2 (2020).

42. Hall, G. et al. Immunohistochemistry for PMS2 and MSH6 alone can replace a four antibody panel for mismatch repair deficiency screening in colorectal adenocarcinoma. Pathology 42, 409–413, doi:10.3109/00313025.2010.493871 (2010).

43. Ijsselsteijn, M. E., van der Breggen, R., Farina Sarasqueta, A., Koning, F. & de Miranda, N. F. C. C. A 40-Marker Panel for High Dimensional Characterization of Cancer Immune Microenvironments by Imaging Mass Cytometry. Frontiers in immunology 10, 2534–2534, doi:10.3389/fimmu.2019.02534 (2019).

44. Berg, S. et al. ilastik: interactive machine learning for (bio)image analysis. Nat Methods 16, 1226–1232, doi:10.1038/s41592-019-0582-9 (2019).

45. Ijsselsteijn, M. E., Somarakis, A., Lelieveldt, B. P. F., Höllt, T. & de Miranda, N. Semi-automated background removal limits data loss and normalises imaging mass cytometry data. Cytometry A, doi:10.1002/cyto.a.24480 (2021).

46. Carpenter, A. E. et al. CellProfiler: image analysis software for identifying and quantifying cell phenotypes. Genome Biol 7, R100, doi:10.1186/gb-2006-7-10-r100 (2006).

47. Somarakis, A., Van Unen, V., Koning, F., Lelieveldt, B. P. F. & Hollt, T. ImaCytE: Visual Exploration of Cellular Microenvironments for Imaging Mass Cytometry Data. IEEE transactions on visualization and computer graphics, 10.1109/TVCG.2019.2931299, doi:10.1109/TVCG.2019.2931299 (2019).

48. van der Maaten, L. J. P. & Hinton, G. E. Visualizing high-dimensional data using t-SNE. J. Mach. Learn. Res. 9, 2579–2605 (2008).

49. Höllt, T. et al. Cytosplore: Interactive Immune Cell Phenotyping for Large Single-Cell Datasets. 35, 171–180, doi:https://doi.org/10.1111/cgf.12893 (2016).

50. Stoeckius, M. et al. Simultaneous epitope and transcriptome measurement in single cells. Nat Methods 14, 865–868, doi:10.1038/nmeth.4380 (2017).

51. Stuart, T. et al. Comprehensive Integration of Single-Cell Data. Cell 177, 1888–1902 e1821, doi:10.1016/j.cell.2019.05.031 (2019).

52. McGinnis, C. S. et al. MULTI-seq: sample multiplexing for single-cell RNA sequencing using lipid-tagged indices. Nat Methods 16, 619–626, doi:10.1038/s41592-019-0433-8 (2019).

53. Haghverdi, L., Lun, A. T. L., Morgan, M. D. & Marioni, J. C. Batch effects in single-cell RNA-sequencing data are corrected by matching mutual nearest neighbors. Nat Biotechnol 36, 421–427, doi:10.1038/nbt.4091 (2018).

54. McInnes, L., Healy, J. & Melville, J. J. a. p. a. Umap: Uniform manifold approximation and projection for dimension reduction. (2018).

55. Dijkstra, K. K. et al. Generation of Tumor-Reactive T Cells by Co-culture of Peripheral Blood Lymphocytes and Tumor Organoids. Cell 174, 1586–1598.e1512, doi:10.1016/j.cell.2018.07.009 (2018).

56. Cattaneo, C. M. et al. Tumor organoid-T-cell coculture systems. Nat Protoc 15, 15–39, doi:10.1038/s41596-019-0232-9 (2020).

57. Dutta, I., Postovit, L. M. & Siegers, G. M. Apoptosis Induced via Gamma Delta T Cell Antigen Receptor “Blocking” Antibodies: A Cautionary Tale. Front Immunol 8, 776, doi:10.3389/fimmu.2017.00776 (2017).

